# An empirical evaluation of functional alignment using inter-subject decoding

**DOI:** 10.1101/2020.12.07.415000

**Authors:** Thomas Bazeille, Elizabeth DuPre, Hugo Richard, Jean-Baptiste Poline, Bertrand Thirion

## Abstract

Inter-individual variability in the functional organization of the brain presents a major obstacle to identifying generalizable neural coding principles. Functional alignment—a class of methods that matches subjects’ neural signals based on their functional similarity—is a promising strategy for addressing this variability. To date, however, a range of functional alignment methods have been proposed and their relative performance is still unclear. In this work, we benchmark five functional alignment methods for inter-subject decoding on four publicly available datasets. Specifically, we consider three existing methods: piecewise Procrustes, searchlight Procrustes, and piecewise Optimal Transport. We also introduce and benchmark two new extensions of functional alignment methods: piecewise Shared Response Modelling (SRM), and intra-subject alignment. We find that functional alignment generally improves inter-subject decoding accuracy though the best performing method depends on the research context. Specifically, SRM and Optimal Transport perform well at both the region-of-interest level of analysis as well as at the whole-brain scale when aggregated through a piecewise scheme. We also benchmark the computational efficiency of each of the surveyed methods, providing insight into their usability and scalability. Taking inter-subject decoding accuracy as a quantification of inter-subject similarity, our results support the use of functional alignment to improve inter-subject comparisons in the face of variable structure-function organization. We provide open implementations of all methods used.

## 1. Introduction

A core challenge for cognitive neuroscience is to find similarity across neural diversity (Churchland 1998); that is, to find shared or similar neural processes supporting the diversity of individual cognitive experience. Anatomical variability and limited structure-function correspondence across cortex (Paquola et al. 2019, Vázquez-Rodríguez et al. 2019) make this goal challenging (Rademacher et al. 1993, Thirion et al. 2006). Even after state-of-the-art anatomical normalization to a standard space, we still observe differences in individual-level functional activation patterns that hinder cross-subject comparisons (Langs et al. 2010, Sabuncu et al. 2010). With standard processing pipelines, it is therefore difficult to disentangle whether individuals are engaging in idiosyncratic cognitive experience *or* if they are engaging in shared functional states that are differently encoded in the supporting cortical anatomy.

To address this challenge, *functional alignment* is an increasingly popular family of methods for functional magnetic resonance imaging (fMRI) analysis: from the initial introduction of hyperalignment in Haxby et al. 2011, the range of associated methods has grown to include Shared Response Modelling (SRM; Chen et al. 2015) and Optimal Transport (Bazeille et al. 2019) with many variations thereof (see e.g. Xu et al. 2018, Yousefnezhad and Zhang 2017, among others). Although this class of methods is broadly referred to as both *functional alignment methods* and *hyperalignment methods*, we adopt the term *functional alignment methods* to better distinguish from the specific Procrustes-based hyper-alignment implementation in use in the literature.

The conceptual shift from anatomically-based to functionally-driven alignment has opened new avenues for exploring neural similarity and diversity. In particular, by aligning activation patterns in a high-dimensional functional space (i.e., where each dimension corresponds to a voxel), we can discover shared representations that show similar trajectories in functional space but rely on unique combinations of voxels across subjects. For a review of current applications of functional alignment, see Haxby et al. 2020.

Nonetheless, it remains unclear how researchers should choose among the available functional alignment methods for a given research application. We therefore aimed to benchmark performance of existing functional alignment methods on several publicly accessible fMRI datasets, with the goal of systematically evaluating their usage for a range of research questions. We consider performance to include both (1) improving inter-subject similarity while retaining individual signal structure as well as (2) computational efficiency, as the latter is an important consideration for scientists who may not have access to specialized hardware. Here, we specifically focus on pair-wise alignments wherein subjects are directly aligned to a target subject’s functional activations. An alternative approach is known as template-based alignment, wherein a group-level functional template is first created and then used as a reference space to which individual functional activations are aligned. Although template-based approaches are an important area of research—particularly for datasets with a large number of subjects—the question of how best to generate the reference template is distinct from its alignment and beyond the scope of the current work. For all alignment methods considered here, technically up-to-date and efficient implementations to reproduce these results are provided at https://github.com/neurodatascience/fmralign-benchmark.

### 1.1. Defining levels of analysis: region-of-interest or whole-brain

Functionally aligning whole-brain response patterns at the voxel level is computationally demanding and may yield biologically implausible transformations (e.g., aligning contralateral regions). Therefore, currently available functional alignment methods generally define transformations within a sub-region. This constraint acts as a form of regularization, considering local inter-subject variability rather than global changes such as large-scale functional reorganization. It also divides the computationally intractable problem of matching the whole-brain into smaller, more tractable sub-problems.

An important consideration, then, is how to define a local neighborhood. Broadly, two main strategies exist: (1) considering voxels within a given region of interest (ROI) that reflects prior expectations on the predictive pattern or (2) grouping or parcellating voxels into a collection of subregions across the whole-brain. Existing functional alignment methods have been proposed using both approaches. For example, the initial introduction of hyperalignment in Haxby et al. 2011 was evaluated within a ventral temporal cortex ROI and was later extended to aggregate many local alignments into larger transforms using a *Searchlight* scheme (Guntupalli et al. 2016). Other methods such as Optimal Transport have been evaluated on whole-brain parcellations (Bazeille et al. 2019), where transforms are derived for each parcel in parallel and then aggregated into a single whole-brain transform. Throughout this work, we therefore consider functional alignment methods at both the ROI and aggregated whole-brain level of analysis.

### 1.2. Quantifying the accuracy of functional alignment

#### 1.2.1. Image-based statistics

A key question is how to objectively measure the performance of functional alignment. One approach is to consider alignment as a reconstruction problem, where we aim to learn a functional alignment transformation that allows us to impute missing images in a target subject using data from source subjects. These functionally aligned maps can then be compared with held-out ground-truth maps from the target subject. We can quantify this comparison using image-based statistics such as the correlation of voxel activity profiles across tasks (Guntupalli et al. 2016, Jiahui et al. 2020), spatial correlation or Dice coefficient between estimated and held-out brain maps (Langs et al. 2014) or other metrics such as *reconstruction ratio* (Bazeille et al. 2019). However, these image-based statistics are sensitive to low-level image characteristics (e.g., smoothness, scaling), and their values can therefore reflect trivial image processing effects (such as the smoothness introduced by resampling routines) rather than meaningful activity patterns.

#### 1.2.2. Adopting a predictive framework to quantify alignment accuracy

Rather than using image-based statistics, an alternative approach is to test functional alignment accuracy in a predictive framework. Prior work adopting this framework has used tests such as time-segment matching from held-out naturalistic data (e.g., Chen et al. 2015, Guntupalli et al. 2016). However, because time-segment matching relies on the same stimulus class to train and test the alignment, it is unclear whether the learnt functional transformations extend to other, unrelated tasks— particularly tasks with low inter-subject correlation (Nastase et al. 2019). We are therefore specifically interested in predictive frameworks that probe model validity by measuring accuracy on held-out data from a different stimulus class, with or without functional alignment.

Inter-subject decoding is a well-known problem in the literature aimed at uncovering generalizable neural coding principles. More in detail, in inter-subject decoding one learns a predictive model on a set of subjects and then test that model on held-out subjects, measuring the extent to which learned representations generalize across individuals. In an information-mapping framework (Kriegeskorte and Diedrichsen 2019), decoding allows one to assess the mutual information between task conditions. Alternate information-mapping approaches include Representational Similarity Analysis (Kriegeskorte et al. 2008), which assesses similarities between relative patterns of activations across task conditions. In this context, functional alignment should facilitate information-mapping by increasing the similarity of condition-specific representations across subjects, thus improving their decoding.

Although the link between mutual information and decoding accuracy is non-trivial (Olivetti et al. 2011), we consider that measuring alignment with decoding accuracy on unseen subjects better fulfils neuroscientists’ expectations of inter-subject alignment in two main ways. First, decoding accuracy provides a more interpretable assessment of performance than other measures such as mutual information estimates. Second, decoding accuracy on a held-out sample provides insight into the external validity and therefore generalizability of derived neural coding principles. Compared to image-based measures, decoding accuracy is thus a more rigorous measure of whether functional alignment improves the similarity of brain signals across subjects while also preserving their structure and usability for broader research use cases. In this work, we therefore quantify functional alignment accuracy by assessing improvements in inter-subject decoding when using functional alignment over and above anatomical alignment. That is, the field-standard approach of normalizing subjects to a standardized anatomical template using diffeomorphic registrations, as implemented in e.g. fMRIPrep (Esteban et al. 2019).

### 1.3. The present study

Using this inter-subject decoding framework, we: (1) establish that functional alignment improves decoding accuracy above anatomical-only alignment, (2) investigate the impact of common methodological choices such as whether alignment is learned in subregions across the whole brain or in a pre-defined region-of-interest (ROI), and (3) compare the impact of specific alignment methods in whole-brain and ROI-based settings. We then provide a qualitative comparison of the transformations learnt by each method to “open the black box” and provide insights into how potential accuracy gains are achieved. Finally, we discuss the availability, usability and scalability of current implementations for each of the methods considered.

## 2. Materials and Methods

In this section, we first consider frameworks for aggregating local functional alignment transformations into a single, larger transform (Section 2.1.1) that can be applied at a whole-brain scale. We then proceed by introducing mathematical notations for functional alignment, as well as the alignment methods included in our benchmark (Section 2.2). We next describe our procedure to quantify alignment performance using inter-subject decoding (Section 2.3) and a series of experiments aimed at investigating the impact of functional alignment on decoding accuracy (Section 2.4). Finally, we describe the datasets (Section 2.5) and implementations used to run each experiment (Section 2.6).

### 2.1. Aggregating local alignments

#### 2.1.1. Comparing searchlight and piecewise schemes

As discussed in Section 1.1, alignment methods are closely linked with the definition of local correspondence models. To align the entire cortex across subjects, two main frameworks have been proposed: searchlight and piecewise aggregation schemes. Each of these frameworks use functional alignment methods to learn local transformations and aggregate them into a single largescale alignment; however, searchlight and piecewise differ in how they aggregate transforms, as illustrated in Figure 2. The *searchlight* scheme (Kriegeskorte et al. 2006), popular in brain imaging (Guntupalli et al. 2018, 2016), has been used as a way to divide the cortex into small overlapping spheres of a fixed radius. This method allows researchers to remain agnostic as to the location of functional or anatomical boundaries, such as those suggested by parcellation-based approaches. A local transform can then be learnt in each sphere and the full alignment is obtained by aggregating (e.g. summing as in Guntupalli et al. 2016 or averaging) across overlapping transforms. Importantly, the aggregated transformation produced is no longer guaranteed to bear the type of regularity (e.g orthogonality, isometry, or diffeomorphicity) enforced during the local neighborhood fit.

**FIG. 1.**
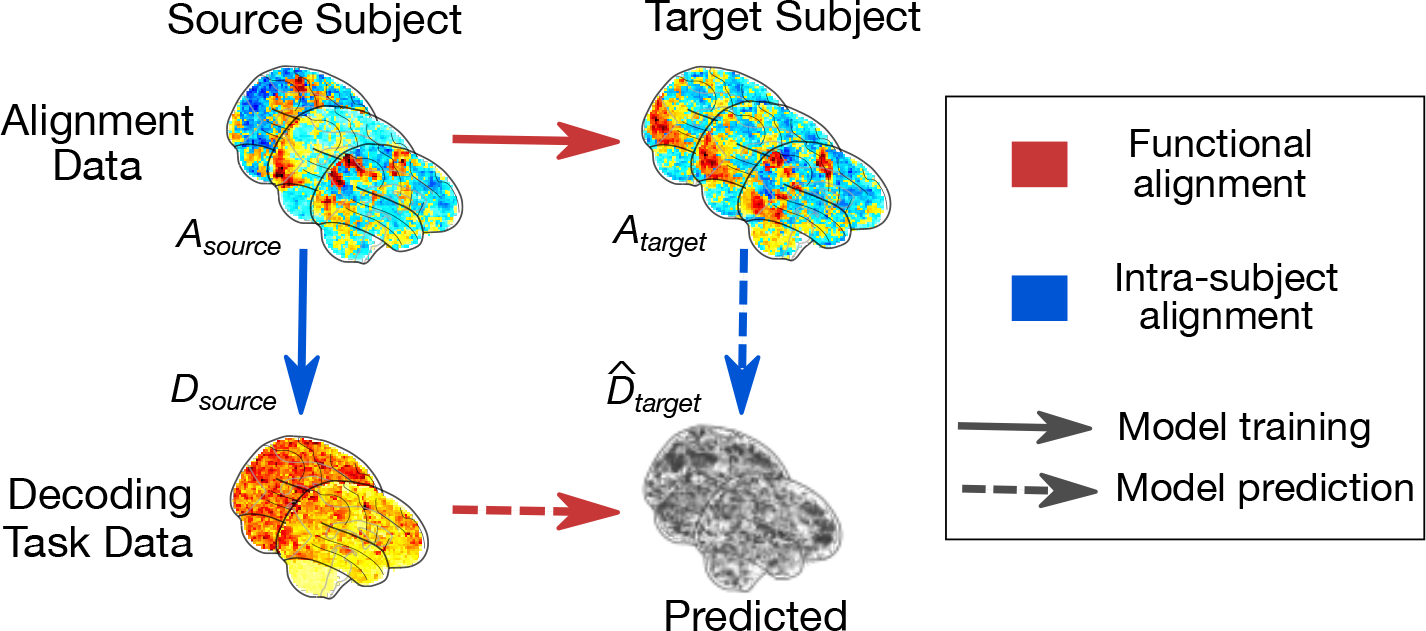
Principle of functional alignment. The goal of functional alignment is to learn correspondence between data drawn from two subjects: from a **source** subject to a **target** subject using their synchronized **alignment** data **A**. In this paper, each subject comes with additional **decoding** task data **D**. Red arrows describe functional alignment methods where correspondence is learnt from **A***^source^* to **A***^target^*, while blue arrow describes intra-subject alignment method, where we learn correlation structure from **A***^source^* to **D***^source^*. Solid arrows indicate a transformation learnt during training. Dashed arrows indicate when the previously learnt transformation is applied in prediction to estimate 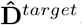.

**FIG. 2.**
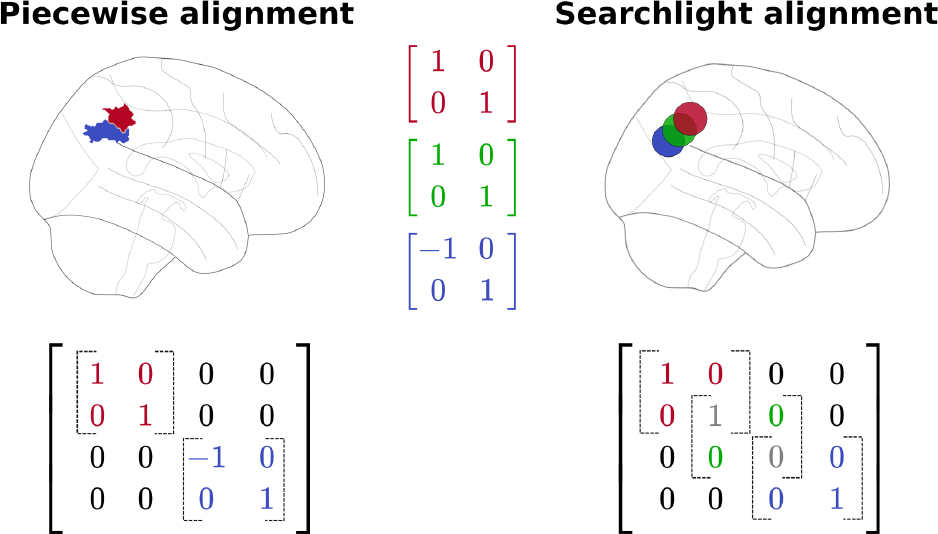
Comparing piecewise and searchlight alignment. In this illustration, transformations are derived for the blue, green, and red areas separately. Note that the piece-wise alignment does not include a green area, as this corresponds to a searchlight overlapping both the red and blue areas. For non-overlapping parcels, these transformations are stacked into a larger orthogonal matrix. For the overlapping searchlight, these transformations are aggregated, with overlapping values averaged. Note that the final transformation for the searchlight alignment is no longer orthogonal in this example.

An alternative scheme, *piecewise alignment* (Bazeille et al. 2019), uses non-overlapping neighborhoods either learnt from the data using a parcellation method—such as k-means—or derived from an *a priori* functional or anatomical atlas. Local transforms are derived in each neighborhood and concatenated to yield a single large-scale transformation. Unlike searchlight, this returns a transformation matrix with the desired regularities. This framework might induce staircase effects or other functionally-irrelevant discontinuities in the final transformation due to the underlying boundaries.

#### 2.1.2. Aggregation schemes used in this benchmark

In the literature to date, searchlight and piecewise aggregation schemes have both been used in conjunction with Generalized Procrustes Analysis (detailed in section 2.2) under the names hyperalignment (Guntupalli et al. 2016) and scaled orthogonal alignment (Bazeille et al. 2019), respectively. We therefore include both search-light Procrustes and piecewise Procrustes in our benchmark. Every other method is regularized at the whole-brain level of analysis through piecewise aggregation.

As piecewise alignment is learnt within a parcellation, an important question is: which brain atlas should be used for piecewise alignment? In Section S4 we compare results from the Schaefer et al. 2018 atlases to those from parcellations derived directly on the alignment data. By default, the results presented below are derived with the 300 ROI parcellation of the Schaefer atlas unless noted otherwise. In the case of searchlight Procrustes, we selected searchlight parameters to match those used in Guntupalli et al. 2016; that is, each searchlight had 5 voxel radius, with a 3 voxel distance between searchlight centers. All searchlight analyses were implemented using PyMVPA (Hanke et al. 2009).

### 2.2. Description of the benchmarked methods

As we use inter-subject decoding to compare functional alignment methods, we only consider methods that meet the following two criteria. First, the alignment transformations should be learnt on activations evoked during temporally synchronized (i.e., co-occuring) task data, or on contrasts matched across individuals. Second, the learnt transformations must be invertible or almost invertible linear mappings and applicable as-is on unseen data with a different task structure. These two criteria exclude several methods currently used in the literature such as regularized canonical correlation analysis (rCCA; Bilenko and Gallant 2016), gradient hyper-alignment (Xu et al. 2018), connectivity hyperalignment (Guntupalli et al. 2018), and methods based on Laplacian embeddings (Langs et al. 2014).

In our whole-brain benchmark, we consider five different alignment methods: searchlight Procrustes (Guntupalli et al. 2016, Haxby et al. 2011), piecewise Procrustes, piecewise Optimal Transport (Bazeille et al. 2019), piecewise Shared Response Modelling (SRM; Chen et al. 2015), and intra-subject correlations across tasks (Tavor et al. 2016), here referred to as “intra-subject alignment.” We provide a brief summary of these methods below.

#### 2.2.1. General notations

Assume that for every subject we have alignment data **A** ∈ ℝ^*p*×*n*^ and decoding task data **D** ∈ ℝ^*p*×*d*^, where *n* is the number of alignment time points or frames, *d* is the number of decoding task images and *p* is the number of voxels. The alignment and decoding task data are collected for both *source* and *target* subjects, which we denote with superscripts.

In general, functional alignment methods learn a transformation matrix **R** ∈ ℝ^*p*×*p*^ that best maps functional signals from a source subject to those of a target subject. To do so, **R** can be seen as a linear mixing of *source* voxels signals such that **RA***^source^* best matches **A***^target^*. **R** is then applied on separate, held-out data from the source subject, **D***^source^* to estimate **D***^target^*. Because we are only learning an estimate of that held-out decoding task data, we denote this 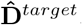. Thus, 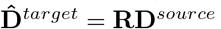.

We consider one method, intra-subject alignment, which uses the same alignment and decoding task data to learn a different transformation than the one described above. Specifically, in intra-subject alignment we are interested in learning **R***^intra^* ∈ ℝ^*n*×*s*^; that is, the “intra-subject” correlations between **A***^source^* and **D***^source^*. We can then use **R***^intra^* to output 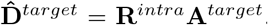. Thus, the main distinction here is that intra-subject alignment does not learn a source-target mapping; instead, it learns a **A** to **D** mapping within-subjects. These notations are illustrated in Figure 1.

#### 2.2.2. Procrustes

Generalized Procrustes analysis, introduced to the cognitive neuroscience literature as *hyperalignment* (Haxby et al. 2011), searches for an orthogonal local transformation **R** to align subject-level activation patterns such that:

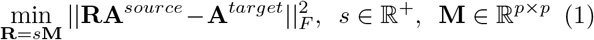

where *p* is the number of voxels in a given region, such that

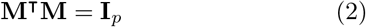

This transform can be seen as a rotation matrix mixing signals of voxels in **A^source^** to reconstruct the signal of voxels in **A^target^** as accurately as possible. We note that hyperalignment as defined in (Haxby et al., 2011) uses a three stage alignment-and-averaging procedure to extend these Procrustes transformations into a group-level, template-based method. In the context of pairwise alignments, however, this method is naturally equivalent to Procrustes. Thus, in the rest of this work we use the terms “hyperalignment” and “Procrustes” interchangeably. As described in Section 2.1.2, we compare two whole-brain implementations of this method: piece-wise Procrustes and searchlight Procrustes, that differ in the way local transformations are aggregated.

#### 2.2.3. Optimal Transport

Optimal transport—first introduced as a functional alignment method in Bazeille et al. 2019—estimates a local transformation **R** that aligns subject-level activation patterns at a minimal overall cost. Specifically, we can compute the cost of aligning two subject-level activation patterns as Tr(**R · C**), where **C** is the functional dissimilarity—or difference in activation patterns—between source and target, as measured by a pairwise functional distance matrix. Thus, for voxel *i* in **A^source^** and voxel *j* in **A^target^**:

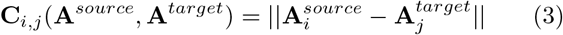

Importantly, the resulting matching is constrained to exhaustively map all source voxels to all target voxels, with every voxel having an equal weight. This implicitly yields an invertible and strongly constrained transform, preserving signal structure as much as possible. To allow for a more efficient estimation, we slightly relax this constraint with an additional entropic smoothing term. As introduced in Cuturi 2013, we can then find **R**, the regularized Optimal Transport plan by finding a minimum for Equation 4 through the Sinkhorn algorithm.

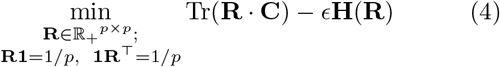

where *ϵ* > 0, and the discrete entropy of the transformation **H**(**R**) is defined as:

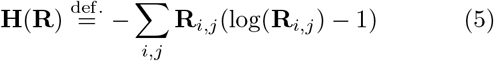

This method differs from Procrustes analysis in that it yields a sparser mapping between source and target voxels with high functional similarity, making it less sensitive to noisy voxels on both ends. The level of sparsity is controlled by *ϵ*, a user-supplied hyper-parameter, which we set to 0.1 throughout our experiments. For our implementation, we rely on the fmralign package. Optimal transport transformations are calculated in a piecewise fashion, following Bazeille et al. 2019.

#### 2.2.4. Shared Response Model

The Shared Response Model (SRM), introduced in Chen et al. 2015, differs from Procrustes analysis and Optimal Transport in that it naturally provides a decomposition of all subjects’s activity at once, rather than requiring pairwise transformations. Specifically, SRM (in its deterministic formulation) estimates a common shared response **S** ∈ ℝ^*k*×*n*^ and a per-subject orthogonal basis **W***^i^* ∈ ℝ^*p*×*k*^ from subject-level alignment data **A***^i^* such that:

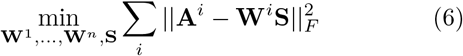

where *n* is the number of time points, *p* is the number of voxels, and *k* is a hyper-parameter indexing the dimensionality. The subject-specific basis **W***^i^* has orthonormal columns such that:

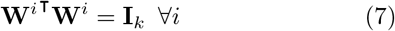

We specifically use the FastSRM implementation proposed by Richard et al. 2019 and available in the BrainIAK library (RRID: SCR 01 4824), that approximates this calculation with an emphasis on improved computational performance. For full details on the computational advantages of FastSRM, we direct the reader to their work.

In order to align our SRM implementation with the other considered alignment algorithms, we introduce a new piecewise SRM method to aggregate SRM transformations across the whole brain. Thus within each parcel or across an *a priori* ROI, SRM decomposes the signal of many subjects in a common basis, with the same orthogonality constraint as Procrustes. This ability to jointly fit inter-subject data through orthogonal transforms makes it reminiscent of Procrustes, with a caveat: SRM is effective if the number of components *k* is large enough to capture all distinct components in the signal.

Given the strong dependency of SRM performance on the selected hyper-parameter *k*, this parameter requires additional experimenter consideration. For piece-wise SRM, we perform a grid search to select the relevant Schaefer parcellation resolution and number of components *k* (see Section S5). From these results, we chose to use Schaefer atlas 700 and run one SRM on each parcel searching for 50 components—or equal to the number of voxels if less than 50 voxels are in a given parcel. For ROI-based analyses, we set *k* to 50 components as in our piecewise analyses and matching the original SRM benchmarks provided in Chen et al. (2015).

#### 2.2.5. Intra-subject alignment

Another alternative to pairwise functional alignment has been proposed in Tavor et al. 2016. In their paper, Tavor and colleagues show that while individual activity patterns in each task may appear idiosyncratic, correspondences learnt across different tasks using a general linear model (e.g., to predict task data from resting-state derived features) display less across-subject variability than individual activity maps. This provides an interesting twist on the typical functional alignment workflow: while most methods learn alignments within a single task and across subjects, we can instead learn within-subject correlations across tasks. The structure of learnt task-specific correlations should then hold in new, unseen subjects. We include here a method for learning these intra-subject correlations in a piecewise fashion, which we call *intra-subject alignment*.

Figure 3 illustrates how we can learn the local-level correlation structure between two independent tasks **A***^source^* ∈ ℝ^*p*×*n*^, **D***^source^* ∈ ℝ^*p*×*d*^ within a single *source* subject. We denote the mapping between these tasks as **R***^intra^* to distinguish it from mappings that are learnt between pairs of subjects.

**FIG. 3.**
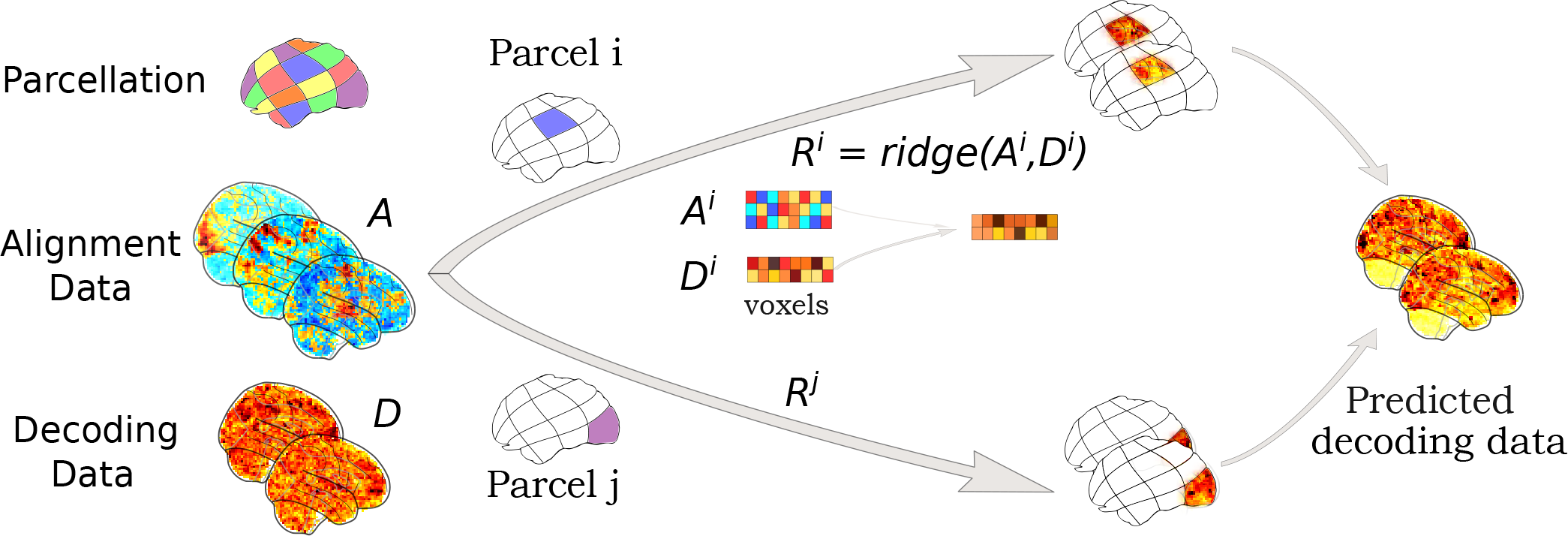
Intra-subject alignment. Using intra-subject alignment to learn piecewise correlations between a single subject’s alignment and decoding task data. As with other piecewise methods, this mapping is learnt separately for all parcels *i … j* of the chosen parcellation. For the *i*th parcel, voxels are samples used to train a cross-validated ridge regression **R***^i^* to map between the two task conditions—alignment data **A_i_** and independent decoding task data **D_i_**—for this source subject. We then aggregate these piecewise predictions into a single, whole-brain prediction 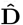. In training, this prediction can be directly compared to the ground-truth decoding data, **D**. When testing, we would have access to the target subject’s alignment data **A** but not their decoding task data, **D**.

From preliminary analyses we observed that—unlike other piecewise techniques (Section S4)—the decoding accuracy for intra-subject alignment strictly improved parcellation resolution so we use the highest resolution Schaefer atlas available (Schaefer et al. 2018). Thus, we first divide alignment and decoding data into 1000 parcels. On a local parcel *i*, each voxel is considered a sample and we train 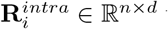 through ridge regression:

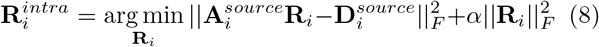

The hyperparameter *α* is chosen with nested cross-validation among five values scaled between 0.1 and 1000 logarithmically.

After repeating this procedure for all *source* subjects, we then use **R***^intra^* to estimate decoding data for *target* subject as 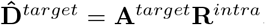. As with other functional alignment methods, we can evaluate the quality of our estimation using an inter-subject decoding framework.

### 2.3. Experimental procedure

For each dataset considered (described in Section 2.5), we calculated the inter-subject decoding accuracy for anatomical-only alignment and for each of the five considered functional alignment methods.

To calculate inter-subject decoding accuracy, we took the trial- or condition-specific beta maps generated for each dataset (see Section 2.5 for full details on beta-map generation) and fit a linear Support Vector Machine (SVM). In order to ensure fair comparisons of decoding accuracy across experiments, we chose a classifier with no feature selection and default model regularization (*C* = 1.0). Classifiers were implemented in scikit-learn (Pedregosa et al. 2011), and decoding accuracy was assessed using a leave-one-subject-out cross-validation scheme. That is, the linear SVM was trained to classify condition labels on all-but-one subject and the resulting trained classifier was used without retraining on the held-out subject, providing an accuracy score for that cross-validation fold.

For each dataset, we first calculated the inter-subject decoding accuracy using anatomical alignment. This served as a baseline accuracy against which we could compare each functional alignment method. Using alignment data, functional alignment transformations were then learnt for each pairwise method, where the left-out subject for that cross-validation fold was the target subject for functional alignment. Inter-subject decoding accuracy was then re-calculated after applying functional alignment transformations to the decoding beta maps.

In the special case of SRM—which allows for calculating an alignment from all provided subjects in a single decomposition—we withheld the left-out subject from the shared response estimation step to avoid data leakage. The projection of the left-out subject is then learnt from previously estimated shared space. Finally, the learnt projections are applied to the decoding data, and decoding is performed on the projected data.

For each cross-validation fold, we report the inter-subject decoding accuracy of a given functional alignment method after subtracting the baseline, anatomical-only accuracy for that same fold. An overview of the experimental procedures is provided in Figure 4.

**FIG. 4.**
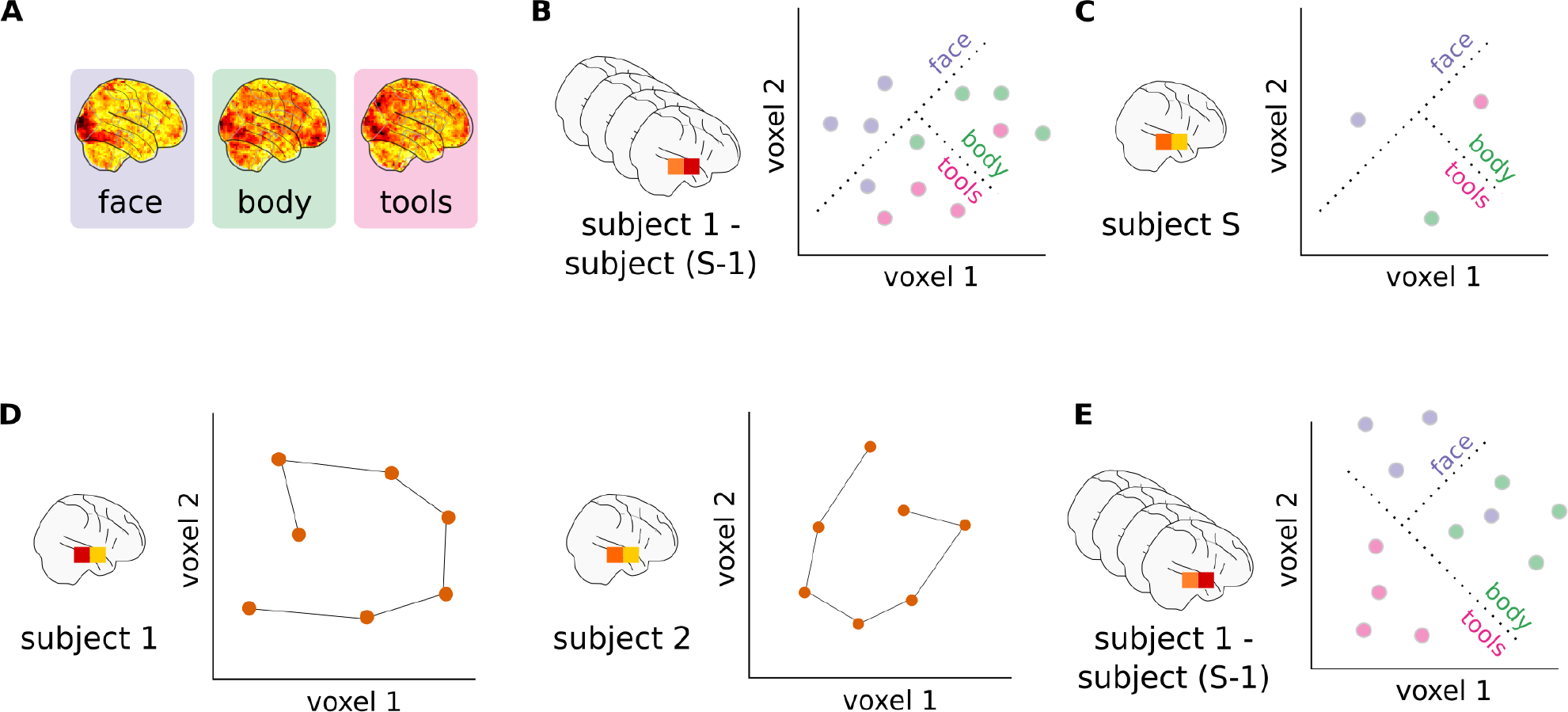
Analysis pipeline. (A) First-level general linear models are fit for each subject to derive trial- or condition-specific beta-maps for each session. (B) These beta maps and their matching condition labels are used to train a linear SVM on the training set of subjects. (C) The trained classifier is applied on a held-out test subject, and accuracy is assessed by comparing the predicted and actual condition labels. (D) On a separate task, we compare subject-level activation patterns as trajectories in the high-dimensional voxel space. This allows us to learn functional alignment transformations that maximize the similarity of these high-dimensional spaces. (E) These voxel-wise transformations are applied on the decoding beta maps, and a new linear SVC is trained to predict condition labels. This trained classifier can then be applied to the held-out test subject and decoding accuracy assessed as in (C).

### 2.4. Main experiments

*Experiment 1* uses the experimental procedure described previously to assess accuracy gains provided by alignment methods with respect to anatomical alignment when applied on whole-brain images. We benchmarked the five methods described in Section 2.2: piecewise Procrustes, searchlight Procrustes, piecewise Optimal Transport, piecewise SRM, and intra-subject alignment, with relevant hyperparameters selected as described previously. Results of this benchmark (on five tasks from four datasets as described in Section 2.5) are presented in Section 3.1. For each method, we also assessed its computation time relative to piecewise Procrustes alignment. Piecewise Procrustes provides a reasonable computational baseline as it is the only considered alignment method that does not include a hyperparameter and therefore shows a stable computation time across experiments.

We estimate the noise ceiling for this task as within-subject decoding accuracy. Within-subject decoding was calculated separately for each subject as the average leave-one-session-out decoding accuracy. We can then directly compare this accuracy value to the inter-subject decoding accuracy when that subject is the target—that is, the left-out—subject. The difference between within- and anatomical inter-subject decoding accuracies, then, is a good approximation of the decoding accuracy lost due to inter-subject variability; therefore, it provides a range of possible accuracy gains that can be expected from functional alignment.

We then conducted *Experiment 2* to understand how whole-brain results compare to ROI-based analyses. Specifically, we replicated *Experiment 1* within selected ROIs, such that local alignment methods were applied directly without any aggregation scheme. ROIs were chosen based on *a priori* expectations of each decoding task (see Section 2.5 for details for each dataset). Results from *Experiment 2* are shown in Section 3.2.

*Experiment 3* tackles the notoriously hard problem of understanding how each of the considered methods align subjects by examining qualitatively their impact on activity patterns across individuals. To “open the black-box,” we reused IBC dataset full-brain alignments learnt in *Experiment 1*. Specifically, we consider the transformation to sub-04’s activity pattern from all other subjects’s functional data. With these transformations, we align two contrasts from each of the two decoding tasks of the IBC dataset: Rapid Serial Visual Presentation of words (RSVP language task) and sound listening. Finally, we run a group conjunction analysis (Heller et al. 2007) on these four aligned contrasts and visualize the results. This statistical analysis, more sensitive than its random effect equivalent on small samples, allows one to infer that every subject activated in the region with a proportion *γ* showing the effect considered. Here we use *γ* = 0.25 to recover all regions selectively activated by at least a few subjects, and we show in Section 3.3 how this group functional topography is modified by alignment.

#### 2.4.1. Control analyses

In addition to our three main experiments, we ran three additional control analyses on the IBC dataset. First, we aimed to assess the impact of the brain parcellation and its resolution on piecewise alignment by comparing whole-brain decoding accuracy for two IBC dataset tasks using piecewise Procrustes across both data-driven and pre-defined parcellations (Section S4). As piecewise SRM displays an interaction between parcellation resolution and the method-specific hyperparameter *k*, we ran an additional grid search for this algorithm to determine its optimal experimental parameters (Section S5).

Second, we calculated inter-subject decoding performance after applying Gaussian smoothing kernels of several widths on both IBC dataset decoding tasks (Section S6). Gaussian smoothing is of particular interest as a comparison to functional alignment, as it is commonly used to facilitate inter-subject comparisons by smoothing over residual variance in functional mappings. Finally, in a third control experiment, we assessed the impact of whether data is represented on the surface or the volume and resolution on decoding accuracy in the IBC RSVP language task (Section S7).

### 2.5. Datasets and preprocessing

In order to assess the performance of each functional alignment method in a range of applications, we searched for publicly accessible datasets that included both a task suitable to learn the alignment (e.g. naturalistic or localizer protocols) as well as an independent decoding task on which we could evaluate functional alignment performance. After discarding datasets where we could not obtain above-chance accuracy levels for within-subject decoding, we retained four datasets: BOLD5000 (Chang et al. 2019), Courtois-NeuroMod (Boyle et al. 2020), Individual Brain Charting (IBC; Pinho et al. 2018), and Study Forrest (Hanke et al. 2016). For the IBC dataset, we included both a language (RSVP language) and auditory (Sounds dataset) decoding task, yielding a total of five decoding tasks that probe visual, auditory and language systems. For a complete description of the alignment and decoding data included in each experiment, please see Table I.

**TABLE I.** Datasets used to benchmark alignment methods. The four datasets used in this benchmark, where each dataset consists of *S* subjects. We note the alignment data used for each dataset and *p* the number of timeframes it comprises. These datasets show the range of possible task structures which work for alignment—from static images for BOLD5000, to statistical contrast maps for IBC, to complex audio or audio-visual movies for Forrest and Courtois Neuromod. A full listing of included 53 contrast maps for IBC is included in Section S8. We also include the decoding task(s) used for each dataset. Each subject’s decoding task data comprises *d* images evenly divided across the listed stimulus categories (except for BOLD5000 categories that are unbalanced). Of note, IBC dataset has two independent decoding tasks, bringing the total number of decoding tasks to five.

| Dataset | $S$ | Alignment data | $p$ | Decoding task | Decoding categories | $d$ |
| --- | --- | --- | --- | --- | --- | --- |
| Individual Brain Charting | 10 | Contrast maps from HCP and ARCHI task batteries | 53 | RSVP Language | Words, Non-Words, Consonants, Sentences, Jabberwocky | 360 |
|  |  |  |  | Sounds dataset | Voice, Nature, Animal, Music, Speech, Tools | 72 |
| BOLD5000 | 4 | COCO, ImageNet, and Scenes images | 300 | Imagenet images content | Plant, Animal, Food, Artifact | 350 |
| Forrest | 10 | *Forrest Gump* audio-movie listening | 1600 | Music genre | Country, Metal, Ambient, Symphonic, Rock | 200 |
| Courtois Neuromod | 6 | *Life* movie watching | 2008 | Visual $n$-back condition | Body 0-back, Body 2-back, Face 0-back, Face 2-back, Place 0-back, Place 2-back, Tools 0-back, Tools 2-back | 72 |

BOLD5000, StudyForrest and Courtois-NeuroMod were preprocessed with fMRIPrep (Esteban et al. 2019), while IBC data were preprocessed using an SPM-based pipeline as described in Pinho et al. 2018. A complete description of the fMRIPrep preprocessing procedures is available in the appendix (Section S1). Preprocessed data were then masked using a grey matter mask, detrended, and standardized using Nilearn (Abraham et al. 2014). To reduce the computational cost of functional alignment, we downsampled all included datasets to 3mm resolution. Both alignment and decoding task data were then additionally smoothed with a 5mm Gaussian kernel. A general linear model (GLM) was fit to each decoding task run to derive trial-specific beta maps (or condition-specific beta maps for the Courtois Neuromod and IBC Sounds tasks), which were carried forward for inter-subject decoding.

As described in Section 2.3, *Experiment 2* uses pre-defined regions of interest (ROIs). We selected large, task-relevant ROIs to ensure that sufficient signal was available when decoding. A large visual region, extracted from the Yeo7 (Buckner et al. 2011) atlas was used for the visual tasks in BOLD5000 and Courtois-NeuroMod. For Forrest and IBC Sounds—which are auditory tasks— we took the Neuroquery (Dockès et al. 2020) predicted response to the term “auditory”. We then compared this predicted response with the BASC (Bootstrap Analysis of Stable Clusters) atlas (at scale 36; Bellec et al. 2010) and took the parcel most overlapping with the predicted response; namely, parcel 25. For IBC RSVP, which is a reading task, we extracted the BASC (at scale 20) atlas components most overlapping with MSDL (Multi-Subject Dictionary Learning; Varoquaux et al. 2011) atlas parcels labeled as left superior temporal sulcus, Broca and left temporo-parietal junction: namely, the 8 and 18 BASC components. We then kept only the largest connected component. All included ROIs are displayed in Figure 7.

### 2.6. Implementation

With the exception of Courtois Neuromod, all other included datasets are available on OpenNeuro (Poldrack et al. 2013) under the following identifiers: *ds000113* (Study Forrest), *ds001499* (BOLD5000), and *ds002685* (IBC). Courtois Neuromod 2020-alpha2 release will be available under a data usage agreement as outlined on https://docs.cneuromod.ca.

Our pipeline entirely relies on open-source Python software, particularly the SciPy stack (Virtanen et al. 2020). All included methods are implemented in fmralign or accessed through their original, open source implementations as described in Section 2.2. To ease replication and extension of the presented results, we have created the `fmralign-benchmark` repository under https://github.com/neurodatascience/fmralign-benchmark. This repository provides an implementation of the procedures adopted in these experiments, building on `fmralign` and previously cited tools.

## 3. Results

### 3.1. Functional alignment improves inter-subject decoding

The *left panel* of Figure 5 displays absolute decoding accuracy change brought by each functional alignment method relative to anatomical alignment on whole-brain images. As every method is trained and tested on the same cross-validation folds, we report the fold-by-fold performance change. The *right panel* displays each method’s relative computation time compared to piecewise Procrustes alignment. For each panel, each point displayed is the result for one leave-one-subject-out cross validation fold and each color corresponds to one of the five decoding tasks. Note that these timings are based on available implementations — `fmralign` for piecewise alignment methods, `pymvpa2` for searchlight, and `BrainIAK` for SRM— and are therefore subject to change as implementations improve. Nonetheless, these estimates provide insight into the current state-of-the-art.

**FIG. 5.**
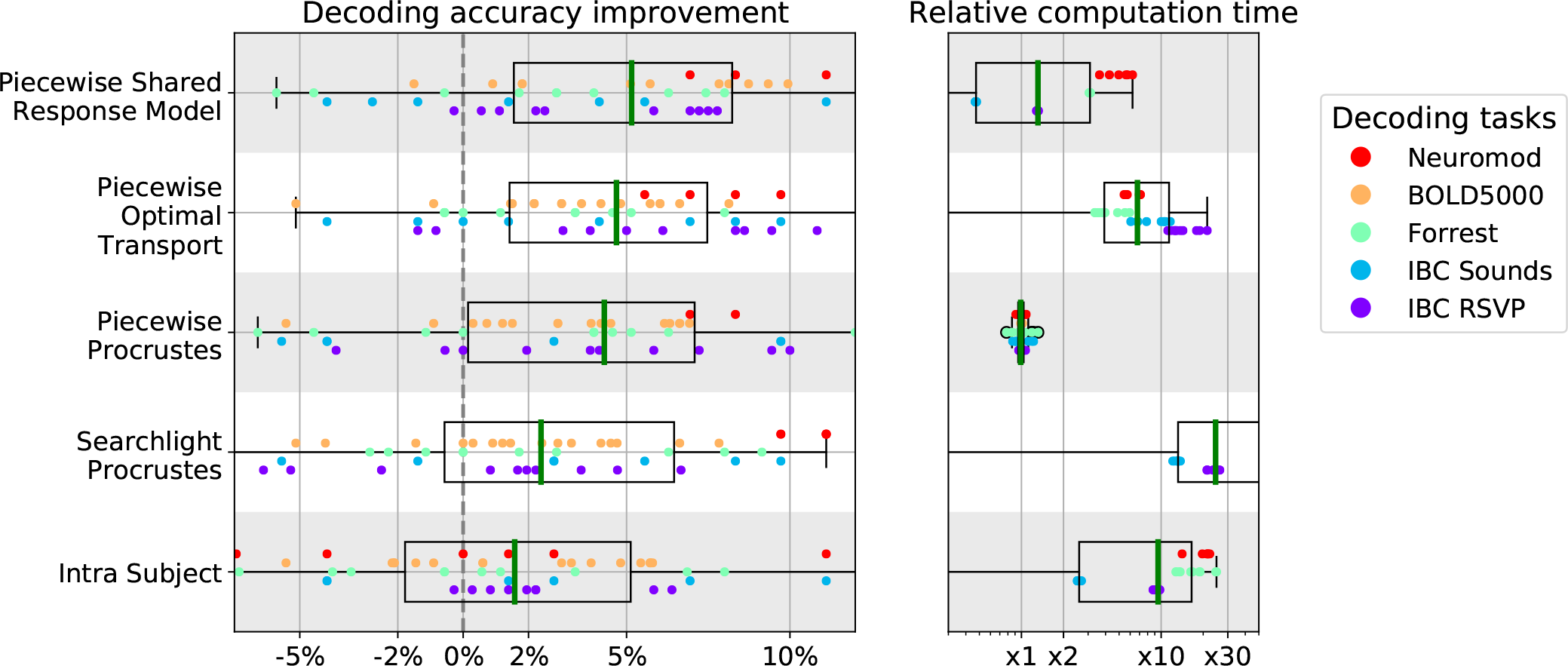
Decoding accuracy improvement and computation time after whole-brain functional alignment. In the *left panel*, we show decoding accuracy improvement for each of the considered functional alignment methods at the whole-brain level of analysis. Each dot represents a single subject, and subjects are colored according to their decoding task. To aggregate results across datasets, we show accuracy scores after subtracting inter-subject decoding accuracy for the same leave-one-subject-out cross-validation fold with anatomical-only alignment. In the *right panel*, we show the computational time for each of the considered methods. All computation times are depicted as relative to piecewise Procrustes. For both panels, each box plot describes the distribution of values across datasets, where the green line indicates the median. All methods seem to improve decoding accuracy across datasets, especially piecewise Shared Response Model, piecewise Optimal Transport and piecewise Procrustes. We also see that piecewise Optimal Transport and searchlight Procrustes are respectively 7 and 25 times slower than piecewise Procrustes.

#### 3.1.1. Alignment substantially improves inter-subject decoding accuracy

Overall, we see that most functional alignment methods consistently improve decoding accuracy, with gains from 2-5% over baseline. This trend is relatively consistent across datasets and target subjects. Thus, alignment methods manage to reliably reduce individual signal variability while preserving task-relevant information in a variety of conditions. Although there is noticeable variance in performance across data sets, these methods generally show significant effects on inter-subject decoding accuracies. As reported in Table S1, baseline accuracy is around 20% above chance on average. In this setting, the observed 5% average improvement across datasets is a substantial increase in performance.

In order to provide further context for these results, we also estimated the noise ceiling for inter-subject decoding. Figure 6 reports that across datasets, the leave-one-session-out (i.e., within-subject) decoding accuracy for the target subject is on average 8.5% higher than the corresponding leave-one-subject-out (i.e., inter-subject) decoding accuracy after anatomical alignment for the same target subject. Thus, we expect that functional alignment methods will achieve at most an 8.5% increase in inter-subject decoding accuracy over anatomical alignment. In this light, we can see that the best functional alignment method recovers more than half of the decoding accuracy lost to inter-subject variability.

**FIG. 6.**
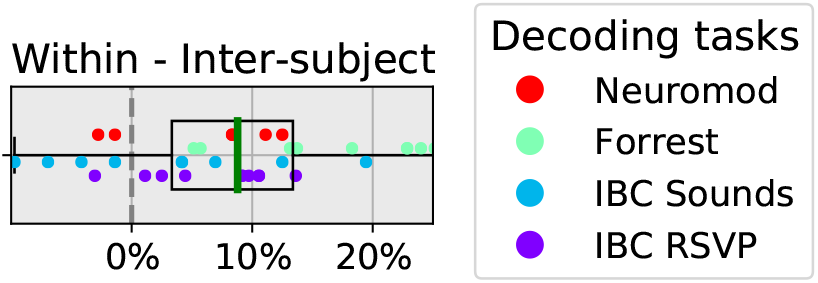
Within-subject minus inter-subject decoding accuracy. We show the difference between the average leave-one-session-out within-subject decoding accuracy and anatomically-aligned leave-one-subject-out inter-subject decoding accuracy, when that target subject is left-out. Thus, each dot corresponds to a single subject, and the dot’s color indicates the decoding task. Of note, BOLD5000 was dropped as it did not have independent folds, and therefore could not be used for within-subject cross-validation. The box plot describes the distribution of differences, where the green line represents the median value. Considering that this difference approximates the effects of inter-individual variability, the best average accuracy improvement one can hope for using functional alignment is around 9%.

Additional control analyses suggest that this effect cannot be explained by smoothing (Section S6). We further find that the presented results are largely insensitive both to whether the data is represented on the cortical surface or in volumetric space as well as to the parcellation resolution used (see section S7).

#### 3.1.2. Piecewise methods show computational and accuracy advantages

Procrustes alignment results in better inter-subject decoding accuracies when performed in a piecewise as compared to a searchlight approach. Specifically, searchlight Procrustes shows lower decoding accuracies on average, suggesting that its internal averaging destroys part of the signal structure recovered by Procrustes. With respect to computational cost, we can see that searchlight Procrustes is 25 times slower on average than piecewise Procrustes. These results suggest that piecewise alignment is a better choice when calculating functional alignment transformations on full-brain data. Moreover, Section S4 shows that gains from piecewise alignment are largely insensitive to the resolution and type of parcellation used; i.e., taken from an atlas or learnt directly from subject data.

The two best performing alignment methods also use a piecewise aggregation scheme. Specifically, piecewise SRM and Optimal Transport yield the highest decoding scores, with a slightly lower standard deviation in accuracy scores than Procrustes.

Piecewise SRM is the best performing method and faster to train than piecewise Procrustes for a fixed set of hyperparameters; however, identifying the ideal hyper-parameters for a new dataset requires a computationally costly grid-search. Our results (see Section S5) suggest that, in general, a large number of components *k* and a high-resolution parcellation are likely to give reasonable performance across datasets.

The second best performing method, Optimal Transport, gives non-trivial accuracy gains in most configurations and only rarely decreases decoding accuracy, likely because of the stronger constraints that it imposes. However, this extra-performance comes at a computational cost: it is on average 7 times slower than Procrustes. For data sets without sufficient data or computational power to perform a hyper-parameter grid search for piecewise SRM, we suggest that Optimal Transport offers robust decoding performance with little hyper-parameter tuning. It remains, however, more computationally costly than the reference implementation of piecewise Procrustes.

#### 3.1.3. Task-specific mappings can be learnt within subjects

The intra-subject alignment approach differs from other considered functional alignment methods in that it learns mappings between the alignment data and decoding task data, with the assumption that these mappings can be generalized across subjects. Our results support this assumption, although this method yields gains half as large as the best performing alignment method and comes with a significant computational cost. Part of this cost can be accounted for by the increase in the number of parcels that are used to preserve signal specificity. Nonetheless, using task-specific mappings as a functional alignment method suggests that future work on refining related methods may be a promising direction of research.

### 3.2. Whole-brain alignment outperforms ROI-based alignment

The *left panel* of Figure 7 displays the performance of each functional alignment method relative to anatomical alignment within task-relevant ROIs. The *right panel* displays each method’s relative computation time compared to piecewise Procrustes alignment.

**FIG. 7.**
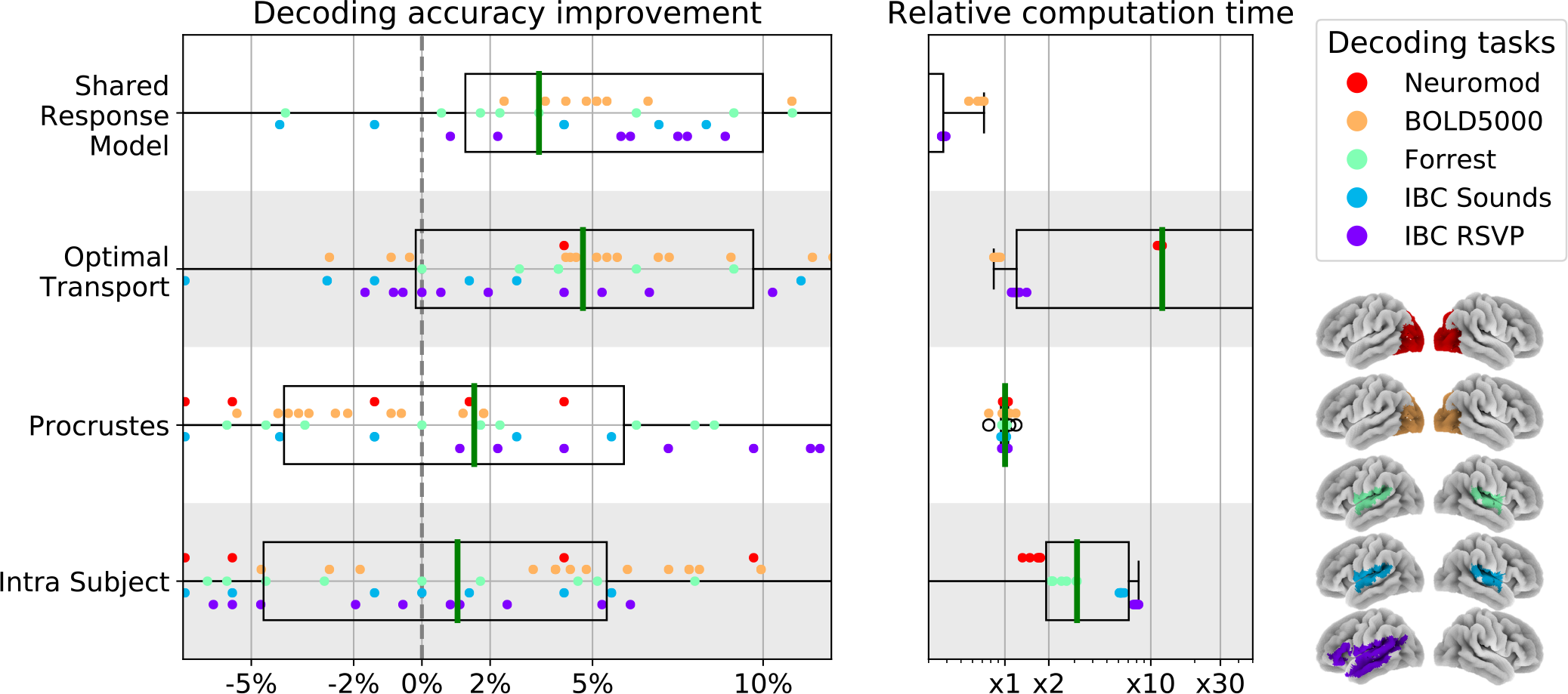
Decoding accuracy improvement and computation time after ROI-based functional alignment. In the *left panel*, we show decoding accuracy for each of the considered local functional alignment methods at the ROI level of analysis. The ROIs used for each dataset are displayed on the *far right*. Each dot represents a single subject, and subjects are colored according to their decoding task. Rather than raw values, we show accuracy scores after subtracting inter-subject decoding accuracy for the same leave-one-subject-out cross-validation fold with anatomical-only alignment. Note that all methods are applied without aggregation, so only the method name is given. In the *right panel*, we show the computational time for each of the considered methods. All computation times are depicted as relative to piecewise Procrustes. For both panels, each box plot describes the distribution of values where the green line indicates the median.

When visually compared to Figure 5, ROI-based decoding accuracies appear to be slightly lower than whole-brain decoding accuracies for most of the methods considered. We directly compare ROI-based and whole-brain alignment in a supplementary analysis, depicted in Figure S1, confirming that ROI-based decoding accuracies are in fact lower on average for the datasets considered in this work. Our results support previous work from the inter-subject decoding literature (Chang et al. 2015, Schrouff et al. 2018) and suggest that full-brain piece-wise alignment yields the best overall decoding pipeline, though we note that this conclusion may change depending on the exact research context.

#### 3.2.1. Optimal Transport and SRM show high ROI performance

Overall, we find that the best performing methods bring a 3-5% improvement in decoding accuracy at the ROI level of analysis. Specifically, Optimal Transport is on average the best performing method, with a median accuracy increase of 5% within task-relevant ROIs. Here, we see that baseline decoding accuracy is less than 10% above chance in all datasets (with the exception of Courtois Neuromod; see Table S2 for exact accuracy values). Thus, the 5% accuracy increase brought by Optimal Transport represents a strong effect.

SRM yields the second best performance within ROIs, showing reasonable decoding accuracy gains on most datasets. It shows more variance across datasets, however, than the other considered methods. In particular, SRM decreases inter-subject decoding accuracy on the visual ROI for Courtois Neuromod, with accuracy values dropping by approximately −20% compared to anatomical alignment (see Table S2). Performance was not significantly improved by using a higher number (up to 600) components, highlighting the unique difficulty in identifying well-suited hyper-parameters for SRM. Interestingly, Procrustes shows substantially lower performance on average in the ROI compared to the whole-brain level of analysis, especially on large ROIs, possibly due to its weak regularization.

Computationally, we see that SRM is the fastest method and runs roughly 3 times faster than Procrustes, while Optimal Transport remains 10 times slower than Procrustes.

We also note that—on average—intra-subject alignment does not show increased inter-subject decoding accuracy within task-relevant ROIs. We suspect that this is likely because when restricting the learnt relationship between data types (e.g. movie-watching to classification task data) to a single ROI, the low number of predicted features precludes the identification of stable multivariate patterns that can transfer across subjects.

### 3.3. Qualitative display of transformations learnt by various methods

Understanding the effects of high-dimensional transformations—such as those used in functional alignment—is non-trivial. To aid in this process, we “open the black box” by functionally aligning a group of subjects to an individual target subject’s functional space and depict the resulting maps in Figure 8. Here, we reuse whole-brain alignments learnt in *Experiment 1*.

**FIG. 8.**
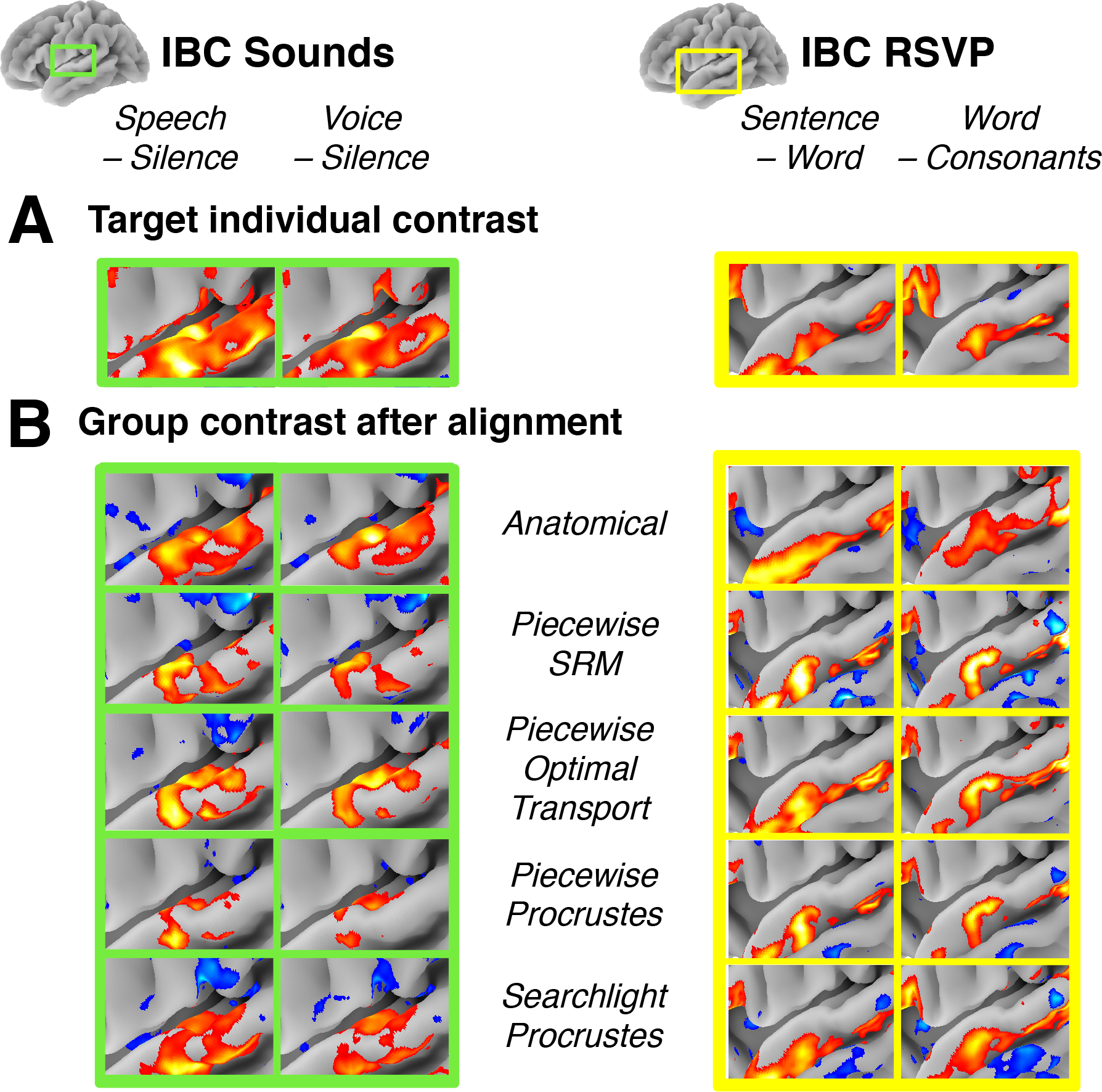
Comparison of alignment methods geometrical effects. (A) Activation patterns for the Target subject (IBC sub-04) for two contrasts from the IBC Sounds task (*Speech* > *Silence, Voice* > *Silence*) and IBC RSVP task (*Sentence* > *Word, Word* > *Consonants*). Here, we only show contrast maps from a sub-region of the temporal lobe containing contrast-relevant information. Note that this sub-region differs slightly between the Sounds and RSVP task. (B) Visualization of a group conjunction analysis of all IBC subjects after alignment to the target subject for each of the considered methods. We used a *γ* value of 0.25 in the group conjunction analysis, which corresponds to at least 25% of the IBC sample showing activation in this temporal region after alignment. For ease of comparison, the colorbar for each contrast and method was scaled to show the full range of values (i.e., the colorbar spans different interval across methods and contrasts) and so is not included here. All displayed maps were thresholded at 1/3 of their maximum value. We see that functional alignment yields stronger contrasts overall when compared to anatomical alignment. Piecewise Procrustes and piecewise Optimal Transport yield less smooth representations, better preserving signal specificity.

We also display the ground-truth individual activation maps in *panel A*, in order to better highlight how each method affects the signal distribution. As a reminder, the contrast data displayed here was not used to learn alignments, so it means that alignment learnt on various task data, not specifically related to language nor audition carried enough information for fine-grain registration of these networks.

We can see that overall, functional alignment methods enhance group-level contrasts compared to anatomical-only alignment; i.e., activation maps are more similar across functionally-aligned subjects. This result is not at the expense of signal specificity, since the aligned group topographies are still sharp. From the comparison between panels *A* and *B*, one can also conclude that alignment methods bring group topography much closer to the targeted subject topography across the considered contrasts. Nonetheless, one can still observe that there seems to be a trade-off between sharpness of activation (low smoothness of image, due to low variance across aligned subjects) with Optimal Transport, and accuracy of their location compared to the target ones (low bias introduced by the matching) with searchlight Procrustes.

## 4. Discussion

In this work, we have proposed a new procedure to measure the information recovered through functional alignment using inter-subject decoding, and we subsequently used this framework to benchmark five functional alignment methods on five distinct decoding tasks across four publicly available datasets.

In general, we find that functional alignment improves inter-subject decoding accuracy in both whole-brain and ROI settings. These results, combined with our qualitative visualization of the effects of functional alignment on signal structure, suggest that functional alignment improves inter-subject correspondence while matching signal to realistic functional topographies. This finding extends and supports conclusions from earlier work (Güçlü and van Gerven 2015, Guntupalli et al. 2016).

At a whole-brain scale, the best performing methods are piecewise SRM, piecewise Optimal Transport, and piecewise Procrustes which each bring 5% improvement over baseline on average. As the baseline inter-subject decoding accuracy is roughly 20% above chance across datasets (Table S1), this 5% increase represents a substantial improvement. We also note that this represents recovering more than half of the accuracy lost to inter-subject variability.

The considered functional alignment methods also improve decoding performance when applied *without* an aggregation scheme (i.e., piecewise or searchlight aggregation) within task-relevant ROIs. Here, Optimal Transport and SRM bring 5% and 3% improvement in inter-subject decoding accuracy, respectively, over a baseline accuracy which is on average 10-15% above chance across datasets (Table S2).

From our control analyses, we observe that these increases in decoding accuracy were reliably greater than the effect of Gaussian smoothing (see section S6). In a minimalistic replication, this effect seems to hold for both volumetric and surface data and at different parcellation resolutions (see section S7; cf. Oosterhof et al. 2011).

Our benchmark also brings new evidence that the latent correspondences that can be learnt between different tasks display less inter-individual variability than the task-specific activation maps (Tavor et al. 2016). *Experiment 1* indeed showed that such correspondences could even be used at a whole-brain scale to transfer signals subjects to solve an inter-subject decoding problem, which is—to the best of our knowledge—an original experimental result. By releasing efficient and accessible implementations of these methods in the fmralign package, we hope to facilitate future cognitive neuroscience research using functional alignment methods.

### 4.1. Combining local alignment models

Across datasets, we find that the aggregation scheme for alignment significantly affects subsequent performance. Notably, piecewise Procrustes outperforms searchlight Procrustes, both in terms of accuracy as well as computational performance. The methodological difference between these aggregation schemes is whether alignment transformations are learnt within overlapping neighborhoods (as in searchlight Procrustes) or not (as in piecewise Procrustes). Searchlight alignment suffers in that the overlap between searchlights requires multiple computations for a given neighborhood, and the aggregated transformation is no longer guaranteed to reflect properties of the original transforms, e.g. orthogonality. Although piecewise aggregation may theoretically introduce discontinuities at parcel boundaries, in our results we do not find evidence of this effect and indeed find that piecewise aggregation overall benefits decoding performance. Importantly, we found that the improved performance of piecewise Procrustes was largely insensitive to parcel size and definition (see Figure S2).

### 4.2. Evaluating alignment performance with decoding

We use inter-subject decoding to quantify the amount of information recovered by functional alignment methods. In general, identifying publicly available datasets with tasks appropriate for both inter-subject decoding as well as functional alignment remains a challenge. Beyond the four datasets included in these results, we investigated several other publicly available datasets such as the Neuroimaging Analysis Replication and Prediction Study (NARPS; Botvinik-Nezer et al. 2020),the Healthy Brain Network Serial Scanning Initiative (HBN-SSI; O’Connor et al. 2017), the interTVA dataset (Aglieri et al. 2019, available as Openneuro *ds001771*) and the Dual Mechanisms of Cognitive Control Project (DMCC, Etzel et al. 2021).

We had difficulties in achieving sufficient baseline accuracy levels in these and other datasets, and we therefore chose not to include them in the present study. This suggests that the amount of signal discriminating complex experimental conditions is not strong enough to find inter-subject patterns robust to variability in many publicly available datasets, likely due to limited sample sizes and unoptimized experimental designs. We hope that broader recognition of the benefits of using inter-subject decoding to uncover neural coding principles across subjects—using functional alignment if necessary—will encourage investigators to collect and share more datasets supporting this type of analysis. Greater data availability will encourage robust, principled comparisons of alignment methods and foster progress in the field.

### 4.3. Study limitations and future directions

Although our study provides a broad evaluation of the performance of several functional alignment methods, there are several dimensions which we hope future work will better address. Notably, we did not thoroughly investigate how alignment performance is impacted by image resolution and whether data are represented on the surface or the volume. Using volumetric images downsampled to a standard resolution of 3mm isotropic enabled us to make fair comparisons across datasets at a reasonable computational cost. We also show in Section S7 that results from piecewise Procrustes alignment on the IBC dataset hold in a higher resolution, surface-based setting. Nonetheless, other functional alignment methods might show different patterns of performance in this setting or at different resolution levels. Moreover, applying these methods on high-resolution images is an exciting perspective to better understand how brain function details vary across subjects. To progress in this direction, a stronger focus on developing computationally efficient methods will be needed. The use of high-resolution parcellations— combined with more efficient implementations of piecewise Optimal Transport or a piecewise Shared Response Model—seem to be particularly promising directions.

We have not examined either the impact of alignment data on the learnt transformations or whether this impact varies across cortex. That is, we could further ask whether certain kinds of stimuli may produce more accurate functional alignments for specialized functional regions. In general, the surveyed functional alignment methods view each subject alignment image as a sample, and the resulting transformation is trained to match corresponding samples across subjects. If some training images lack stable signal in a given ROI, functional alignment methods are unlikely to learn meaningful transformations in this region. Finally this benchmark largely focused on pairwise alignment models. Template-based models—beyond latent factor models as SRM—are an important area of research to further improve the usability of functional alignment methods, particularly in research settings with a large number of subjects. In future work, we intend to address the above questions to learn more about when functional alignment methods are most appropriate.

## 5. Conclusion

In the present work, we have provided an extensive benchmark of five popular functional alignment methods across five unique experimental tasks from four publicly available datasets. Assessing each method in an inter-subject decoding framework, we show that both Shared Response Modelling (SRM) and Optimal Transport perform well at a region-of-interest level of analysis, as well as at the whole-brain scale when aggregated through a piecewise scheme. Our results support previous work proposing functional alignment to improve across-subject comparisons, while providing nuance that some alignment methods may be most appropriate for a given research question. We further suggest that identified improvements in inter-subject decoding demonstrate the potential of functional alignment to identify generalizable neural coding principles across subjects.

## Acknowledgments

This project has received funding from the European Union’s Horizon 2020 Framework Programme for Research and Innovation under the Specific Grant Agreement No. 945539 (Human Brain Project SGA3) and the Digiteo French program. This work was also partially funded by the National Institutes of Health (NIH) NIH-NIBIB P41 EB019936 (ReproNim) NIH-NIMH R01 MH083320 (CANDIShare) and NIH RF1 MH120021 (NIDM), the National Institute Of Mental Health under Award Number R01MH096906 (Neurosynth), as well as the Canada First Research Excellence Fund, awarded to McGill University for the Healthy Brains for Healthy Lives initiative and the Brain Canada Foundation with support from Health Canada.

We wish to thank all researchers that made this study possible by making their datasets publicly available, especially J. Etzel, S. Takerkart, and P. Bellec for kindly taking the time to provide us preprocessed version of their datasets and thorough explanations of their experimental designs. Data from the Courtois project on neural modelling was made possible by a generous donation from the Courtois foundation, administered by the Fondation Institut Gériatrie Montréal at CIUSSS du Centre-Sud-de-l’île-de-Montréal and University of Montreal. The Courtois NeuroMod team is based at Centre de Recherche de l’Institut Universitaire de Gériatrie de Montréal, with several other institutions involved. See the cneuromod documentation for an up-to-date list of contributors (https://docs.cneuromod.ca).

