## Supplementary material for "An empirical evaluation of functional alignment using inter-subject decoding"

### SUPPLEMENTAL MATERIALS

#### S1. FMRIPREP PREPROCESSING

Results included in this manuscript come from preprocessing performed using *fMRIPrep* 20.1.1+38.g8480eabb (Esteban et al. 2018b; Esteban et al. 2018a; RRID:SCR_016216), which is based on *Nipype* 1.5.0 (Gorgolewski et al. 2011; Gorgolewski et al. 2018; RRID:SCR_002502).

##### S1.1. Anatomical data preprocessing

The T1-weighted (T1w) image was corrected for intensity non-uniformity (INU) with N4BiasFieldCorrection (Tustison et al. 2010), distributed with ANTs 2.2.0 (Avants et al. 2008, RRID:SCR_004757), and used as T1w-reference throughout the workflow. The T1w-reference was then skull-stripped with a Nipype implementation of the antsBrainExtraction.sh workflow (from ANTs), using OASIS30ANTs as target template. Brain tissue segmentation of cerebrospinal fluid (CSF), white-matter (WM) and gray-matter (GM) was performed on the brain-extracted T1w using fast (FSL 5.0.9, RRID:SCR_002823, Zhang et al. 2001). Volume-based spatial normalization to one standard space (MNI152NLin2009cAsym) was performed through nonlinear registration with antsRegistration (ANTs 2.2.0), using brain-extracted versions of both T1w reference and the T1w template. The following template was selected for spatial normalization: ICBM 152 Nonlinear Asymmetrical template version 2009c (Fonov et al. 2009, RRID:SCR_008796; TemplateFlow ID: MNI152NLin2009cAsym).

##### S1.2. Functional data preprocessing

For each subject's BOLD runs (across all tasks and sessions), the following preprocessing was performed. First, a reference volume and its skull-stripped version were generated by aligning and averaging 1 single-band references (SBRefs). A B0-nonuniformity map (or *fieldmap*) was estimated based on two (or more) echo-planar imaging (EPI) references with opposing phase-encoding directions, with `3dQwarp` Cox and Hyde (1997) (AFNI 20160207). Based on the estimated susceptibility distortion, a corrected EPI (echo-planar imaging) reference was calculated for a more accurate co-registration with the anatomical reference. The BOLD reference was then co-registered to the T1w reference using `bbregister` (FreeSurfer) which implements boundary-based registration (Greve and Fischl 2009). Co-registration was configured with six degrees of freedom. Head-motion parameters with respect to the BOLD reference (transformation matrices, and six corresponding rotation and translation parameters) are estimated before any spatiotemporal filtering using `mcflirt` (FSL 5.0.9, Jenkinson et al. 2002).

First, a reference volume and its skull-stripped version were generated using a custom methodology of *fMRIPrep*. The BOLD time-series (including slice-timing correction when applied) were resampled onto their original, native space by applying a single, composite transform to correct for head-motion and susceptibility distortions. These resampled BOLD time-series will be referred to as *preprocessed BOLD in original space*, or just *preprocessed BOLD*. The BOLD time-series were resampled into standard space, generating a *preprocessed BOLD run in MNI152NLin2009cAsym space*. First, a reference volume and its skull-stripped version were generated using a custom methodology of *fMRIPrep*.

All resamplings can be performed with *a single interpolation step* by composing all the pertinent transformations (i.e. head-motion transform matrices, susceptibility distortion correction when available, and co-registrations to anatomical and output spaces). Gridded (volumetric) resamplings were performed using `antsApplyTransforms` (ANTs), configured with Lanczos interpolation to minimize the smoothing effects of other kernels (Lanczos 1964).

Many internal operations of *fMRIPrep* use *Nilearn* 0.6.2 (Abraham et al. 2014, RRID:SCR_001362), mostly within the functional processing workflow. For more details of the pipeline, see the section corresponding to workflows in *fMRIPrep*'s documentation.

##### S1.3. Copyright Waiver

The above boilerplate text was automatically generated by fMRIPrep with the express intention that users should copy and paste this text into their manuscripts *unchanged*. It is released under the CC0 license.

### S2. ABSOLUTE DECODING ACCURACY OF VARIOUS METHODS

Tables S1 and S2 report absolute decoding accuracies for *Experiment 1* and *Experiment 2*, to bring a different view of results presented in Figures 5 and 7, as relative improvements brought over anatomical registration by various alignment methods. This "per dataset view" highlight that gains brought by best methods are substantial improvement over baseline, especially when compared to chance.

### S3. WHOLE-BRAIN DECODING PROVIDES BETTER ACCURACY THAN ROI-BASED DECODING

In Figure S1, we compare ROI-based and whole-brain inter-subject decoding accuracy improvements for piecewise Procrustes alignment above anatomical-only alignment. We see that whole-brain alignment generally shows higher inter-subject decoding improvements compared to ROI-based alignment. As mentioned in the main text, this result supports previous work from the inter-subject decoding literature (Chang et al. 2015, Schrouff et al. 2018), and it suggests that full-brain piecewise alignment yields the best overall decoding pipeline.

### S4. PARCELLATION HAS LIMITED IMPACT ON DECODING ACCURACY

To assess the impact of the parcellation used on piecewise alignment results, we compared decoding accuracy gains while varying the parcellation kind and resolution. First, we consider the multi-resolution Schaefer et al. (2018) atlas, which was learnt using a gradient weighted markov random field on resting state data from 1489 subjects. We compare this *a priori* parcellation to two parcellations learnt directly on the subject's alignment data after 5mm FWHM Gaussian smoothing: K-means or Hierarchical K-means. All these parcellations were taken at ten resolutions from 100 to 1000 parcels.

As hierarchical K-means may be less familiar to readers, we briefly describe it in more detail here. This method is a variant of K-means aimed specifically at obtaining more balanced parcels. To identify $k$ parcels, we first apply K-means to cluster the voxels in $\sqrt{k}$ big clusters. Each of these "big clusters" is then clustered again in $\sqrt{k}$ to obtain a total of $k$ smaller well-balanced parcels. In this experiment, K-means and Hierarchical K-means implementations used are respectively from `scikit-learn` and `fmralign`, and fitted as part of `fmralign` alignment functions on the source subject data.

We plot piecewise Procustes accuracy improvements for these three parcellation methods and ten resolutions in Figure S2. Here, we only show the IBC Sounds and IBC RSVP decoding tasks to ease in interpretation. Overall, we observe on these two tasks that the type and resolution of parcellation used does not have a strong impact on accuracy improvements above anatomical-only alignment. We therefore suggest that *piecewise alignment* can be used with confidence that the parcellation choice won't strongly impact its results.

### S5. GRID-SEARCH OF PIECEWISE SRM HYPERPARAMETERS

As a piecewise implementation of SRM is a novel contribution from this work, we had no prior knowledge on how to properly set hyperparameters from this methods (clustering type and resolution as well as the number of components to set for each parcelwise SRM).

For the type of clustering, we limited ourselves to a pre-computed parcellation (the Schaefer atlas) available at various resolutions. This is based on the intuitions acquired on Procrustes ( see Section S4) that the parcellation type was not of utmost importance to decoding results. We ran a cross-validation on the two remaining parameters. We used Schaefer atlas at resolution : [100,300,500,700] while our number of components ranged in [5,25,35,50]. Figure S3 present the results of this cross-validation, that led us to chose Schaefer atlas 700 and 50 components as hyper parameters for our main experiments.

### S6. FUNCTIONAL ALIGNMENT IS NOT MERELY SMOOTHING

Gaussian smoothing is a common preprocessing step in neuroimaging group studies, which reconciles dissimilar subject-level signals by smoothing over inter-individual variability. Our qualitative results (section 3.3) show that best performing alignment methods do not seem to smooth the signal across voxels, but instead preserve the signal specificity while matching its geometry with the target subject functional topography. Specifically, we compared decoding gains from six different Gaussian smoothing kernels to those obtained through the reference method piecewise Procrustes alignment.

The results displayed in Figure S4 clearly support previous findings (Guntupalli et al. 2016) that smoothing does not improve inter-subject decoding performance—and therefore recover mutual information—in the same way as functional alignment.

### S7. IMPACT OF THE DATA REPRESENTATION AND RESOLUTION

Oosterhof et al. 2011 argued that functional alignment benefits from working with a representation of the fMRI signal on the cortical surface (Coalson et al. 2018). Relatedly, we would also expect that the resolution of the data

| Methods/Dataset | IBC RSVP | IBC Sounds | Forrest | BOLD5000 | Neuromod |
|---|---|---|---|---|---|
| **Chance** | 16.7 | 16.7 | 20 | 25.5 | 12.5 |
| **Anatomical** | 38.2 ± 4.1 | 32.7 ±4.8 | 31.4 ± 4.7 | 33.3 ± 2.9 | 54.1 ± 6.7 |
| **Intra-subject** | 39.6 ± 3.1 | 36.4 ± 8.6 | 31.9 ± 5.4 | 34.9 ± 2.8 | 55.6 ± 7.7 |
| **Searchlight Procrustes** | 39.0 ± 5.1 | 32.6 ± 6.5 | **33.6** ± 6.1 | 35.2 ± 2.0 | 65.5 ± 6.6 |
| **Piecewise Procrustes** | 42.0 ± 4.7 | 36.6 ± 5.5 | **33.8** ± 6.4 | 36.4 ± 1.8 | **67.4** ± 10.6 |
| **Piecewise Optimal Transport** | **43.5** ± 5.5 | **38.0** ± 9.5 | **33.8** ± 5.6 | 36.6 ± 2.1 | 65.3 ± 9.1 |
| **Piecewise Shared Response Model** | 42.4 ± 4.0 | 37.0 ± 6.8 | **33.7** ± 7.2 | **39.6** ± 2.9 | 66.2 ± 7.0 |

TABLE S1. **Fullbrain benchmark absolute decoding accuracy** (%)

| Methods/Dataset | IBC RSVP | IBC Sounds | Forrest | BOLD5000 | Neuromod |
|---|---|---|---|---|---|
| **Chance** | 16.7 | 16.7 | 20 | 25.5 | 12.5 |
| **Anatomical** | 22.3 ± 3.3 | 26.9 ± 6.9 | 28.6 ± 6.0 | 33.5 ± 3.3 | 50.2 ± 5.3 |
| **Intra-subject** | 22.0 ± 1.7 | 25.6 ± 2.7 | 27.5 ± 3.2 | 38.0 ± 2.6 | 51.4 ± 10.2 |
| **Procrustes** | **31.0** ± 4.3 | **32.7** ± 8.3 | 30.1 ± 7.0 | 30.1 ± 2.2 | 46.5 ± 7.8 |
| **Optimal Transport** | 24.9 ± 2.5 | 29.9 ± 5.4 | 28.9 ± 2.2 | 39.7 ± 2.7 | **64.4** ± 8.9 |
| **Shared Response Model** | 30.4 ± 4.4 | 30.9 ± 6.3 | **34.3** ± 5.2 | **40.9** ± 3.6 | 32.6 ± 5.5 |

TABLE S2. **ROI benchmark absolute decoding accuracy** (%)

representation—whether in the surface or the volume—will impact the quality of the alignment learnt.

To assess the dependence of our 3mm volumetric results presented in the main text on sampling parameters, we replicated our inter-subject decoding framework with the IBC RSVP language task data on a high-resolution cortical surface representation(*fsaverage7*)(obtained through `freesurfer` surface projection of full-resolution raw images in their respective subject space, later on mapped to the common surface template). This surface mesh includes 168k cortical nodes per hemisphere, which we divided into 350 parcels per hemisphere using *Schaefer* atlas at scale 700.

We provide results for the inter-subject decoding accuracy gains seen with the reference functional alignment method of piecewise Procrustes over standard, anatomical-only alignment. We had to limit to this setting because (i) replicating this analysis on every dataset would represent an important amount of processing work, and (ii) working on other methods than piecewise Procrustes on this very large data is computationally prohibitive.

The results displayed in Figure S5 show that although decoding gains are a little higher using high-resolution surface-based representation, they remain in the same range as the volume-based representation. This shows that a 10-fold higher resolution can help match more precisely topographies across subjects (and reduce the decoding variance as a consequence), but no important marginal gains can be expected from it. In the end the signal available for use is bounded by the same rough limitations: test-retest reliability in each subject.

### S8. IBC ALIGNMENT DATA EXPLAINED

In this work, 53 contrasts were pulled together are used as alignment for IBC dataset. The contrasts are common to all subjects and taken from both HCP and ARCHI protocol. In order they are labelled : *audio left button press, audio right button press, video left button press, video right button press, horizontal checkerboard, vertical checkerboard, audio sentence, video sentence, audio computation, video computation, saccades, rotation hand, rotation side, object grasp, object orientation, mechanistic audio, mechanistic video, triangle mental, triangle random, false belief audio, false belief video, speech sound, non speech sound, face gender, face control, face trusty, expression intention, expression gender, expression control, shape, face, punishment, reward, left hand, right hand, left foot, right foot, tongue, cue, story, math,,relational, match, mental, random, 0back body, 2back body, 0back face, 2back face, 0back tools, 2back tools, 0back place, 2back place.*

To know more, please visit the relevant IBC documentation.

Abraham, A., Pedregosa, F., Eickenberg, M., Gervais, P., Mueller, A., Kossaifi, J., Gramfort, A., Thirion, B., and Varoquaux, G. (2014). Machine learning for neuroimaging with scikit-learn. *Frontiers in Neuroinformatics*, 8.

Avants, B., Epstein, C., Grossman, M., and Gee, J. (2008). Symmetric diffeomorphic image registration with cross-correlation: Evaluating automated labeling of elderly and neurodegenerative brain. *Medical Image Analysis*,

12(1):26–41.

Chang, L. J., Gianaros, P. J., Manuck, S. B., Krishnan, A., and Wager, T. D. (2015). A sensitive and specific neural signature for Picture-Induced negative affect. *PLoS Biol.*, 13(6):e1002180.

Coalson, T. S., Van Essen, D. C., and Glasser, M. F. (2018). The impact of traditional neuroimaging methods on the spatial localization of cortical areas. *Proceedings of the National Academy of Sciences*, 115(27):E6356–E6365.

Cox, R. W. and Hyde, J. S. (1997). Software tools for analysis and visualization of fmri data. *NMR in Biomedicine*, 10(4-5):171–178.

Esteban, O., Blair, R., Markiewicz, C. J., Berleant, S. L., Moodie, C., Ma, F., Isik, A. I., Erramuzpe, A., Kent, James D. andGoncalves, M., DuPre, E., Sitek, K. R., Gomez, D. E. P., Lurie, D. J., Ye, Z., Poldrack, R. A., and Gorgolewski, K. J. (2018a). fmriprep. *Software*.

Esteban, O., Markiewicz, C., Blair, R. W., Moodie, C., Isik, A. I., Erramuzpe Aliaga, A., Kent, J., Goncalves, M., DuPre, E., Snyder, M., Oya, H., Ghosh, S., Wright, J., Durnez, J., Poldrack, R., and Gorgolewski, K. J. (2018b). fMRIPrep: a robust preprocessing pipeline for functional MRI. *Nature Methods*.

Fonov, V., Evans, A., McKinstry, R., Almli, C., and Collins, D. (2009). Unbiased nonlinear average age-appropriate brain templates from birth to adulthood. *NeuroImage*, 47, Supplement 1:S102.

Gorgolewski, K., Burns, C. D., Madison, C., Clark, D., Halchenko, Y. O., Waskom, M. L., and Ghosh, S. (2011). Nipype: a flexible, lightweight and extensible neuroimaging data processing framework in python. *Frontiers in Neuroinformatics*, 5:13.

Gorgolewski, K. J., Esteban, O., Markiewicz, C. J., Ziegler, E., Ellis, D. G., Notter, M. P., Jarecka, D., Johnson, H., Burns, C., Manhães-Savio, A., Hamalainen, C., Yvernault, B., Salo, T., Jordan, K., Goncalves, M., Waskom, M., Clark, D., Wong, J., Loney, F., Modat, M., Dewey, B. E., Madison, C., Visconti di Oleggio Castello, M., Clark, M. G., Dayan, M., Clark, D., Keshavan, A., Pinsard, B., Gramfort, A., Berleant, S., Nielson, D. M., Bougacha, S., Varoquaux, G., Cipollini, B., Markello, R., Rokem, A., Moloney, B., Halchenko, Y. O., Wassermann, D., Hanke, M., Horea, C., Kaczmarzyk, J., de Hollander, G., DuPre, E., Gillman, A., Mordom, D., Buchanan, C., Tungaraza, R., Pauli, W. M., Iqbal, S., Sikka, S., Mancini, M., Schwartz, Y., Malone, I. B., Dubois, M., Frohlich, C., Welch, D., Forbes, J., Kent, J., Watanabe, A., Cumba, C., Huntenburg, J. M., Kastman, E., Nichols, B. N., Eshaghi, A., Ginsburg, D., Schaefer, A., Acland, B., Giavasis, S., Kleesiek, J., Erickson, D., Küttner, R., Haselgrove, C., Correa, C., Ghayoor, A., Liem, F., Millman, J., Haehn, D., Lai, J., Zhou, D., Blair, R., Glatard, T., Renfro, M., Liu, S., Kahn, A. E., Pérez-García, F., Triplett, W., Lampe, L., Stadler, J., Kong, X.-Z., Hallquist, M., Chetverikov, A., Salvatore, J., Park, A., Poldrack, R., Craddock, R. C., Inati, S., Hinds, O., Cooper, G., Perkins, L. N., Marina, A., Mattfeld, A., Noel, M., Snoek, L., Matsubara, K., Cheung, B., Rothmei, S., Urchs, S., Durnez, J., Mertz, F., Geisler, D., Floren, A., Gerhard, S., Sharp, P., Molina-Romero, M., Weinstein, A., Broderick, W., Saase, V., Andberg, S. K., Harms, R., Schlamp, K., Arias, J., Papadopoulos Orfanos, D., Tarbert, C., Tambini, A., De La Vega, A., Nickson, T., Brett, M., Falkiewicz, M., Podranski, K., Linkersdörfer, J., Flandin, G., Ort, E., Shachnev, D., McNamee, D., Davison, A., Varada, J., Schwabacher, I., Pellman, J., Perez-Guevara, M., Khanuja, R., Pannetier, N., McDermottroe, C., and Ghosh, S. (2018). Nipype. *Software*.

Greve, D. N. and Fischl, B. (2009). Accurate and robust brain image alignment using boundary-based registration. *NeuroImage*, 48(1):63–72.

Guntupalli, J. S., Hanke, M., Halchenko, Y. O., Connolly, A. C., Ramadge, P. J., and Haxby, J. V. (2016). A model of representational spaces in human cortex. *Cereb. Cortex*, 26(6):2919–2934.

Jenkinson, M., Bannister, P., Brady, M., and Smith, S. (2002). Improved optimization for the robust and accurate linear registration and motion correction of brain images. *NeuroImage*, 17(2):825–841.

Lanczos, C. (1964). Evaluation of noisy data. *Journal of the Society for Industrial and Applied Mathematics Series B Numerical Analysis*, 1(1):76–85.

Oosterhof, N. N., Wiestler, T., Downing, P. E., and Diedrichsen, J. (2011). A comparison of volume-based and surface-based multi-voxel pattern analysis. *Neuroimage*, 56(2):593–600.

Schaefer, A., Kong, R., Gordon, E. M., Laumann, T. O., Zuo, X.-N., Holmes, A. J., Eickhoff, S. B., and Yeo, B. T. T. (2018). Local-Global parcellation of the human cerebral cortex from intrinsic functional connectivity MRI. *Cereb. Cortex*, 28(9):3095–3114.

Schrouff, J., Monteiro, J. M., Portugal, L., Rosa, M. J., Phillips, C., and Mourao-Miranda, J. (2018). Embedding Anatomical or Functional Knowledge in Whole-Brain Multiple Kernel Learning Models. *Neuroinformatics*, 16(1):117–143.

Tustison, N. J., Avants, B. B., Cook, P. A., Zheng, Y., Egan, A., Yushkevich, P. A., and Gee, J. C. (2010). N4itk: Improved n3 bias correction. *IEEE Transactions on Medical Imaging*, 29(6):1310–1320.

Zhang, Y., Brady, M., and Smith, S. (2001). Segmentation of brain MR images through a hidden markov random field model and the expectation-maximization algorithm. *IEEE Transactions on Medical Imaging*, 20(1):45–57.

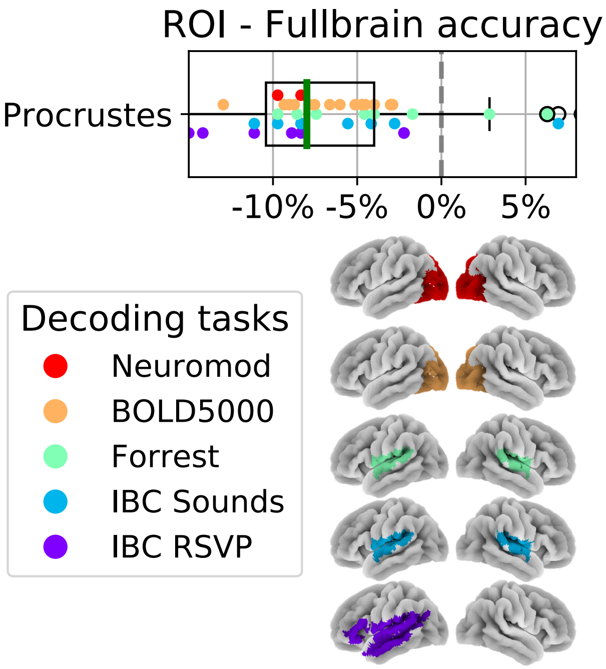


FIG. S1. **Comparing ROI and whole-brain decoding accuracy after piecewise Procrustes alignment.** The ROIs used for each dataset are displayed on the *lower panel*. In the *upper panel*, we show the distribution of differences in decoding accuracy scores between ROI-based and whole-brain piecewise Procrustes alignment. Each dot represents a single subject, and subjects are colored according to their decoding task. Each difference score is calculated by subtracting the inter-subject decoding accuracy for whole-brain piecewise Procrustes alignment from the ROI-based piecewise Procrustes alignment accuracy score—for the same leave-one-subject-out cross-validation fold. The box plot thus describes the distribution of differences, where the green line represents the median value. We see that decoding accuracy is lower when performed within ROIs than when performed on the whole-brain data.

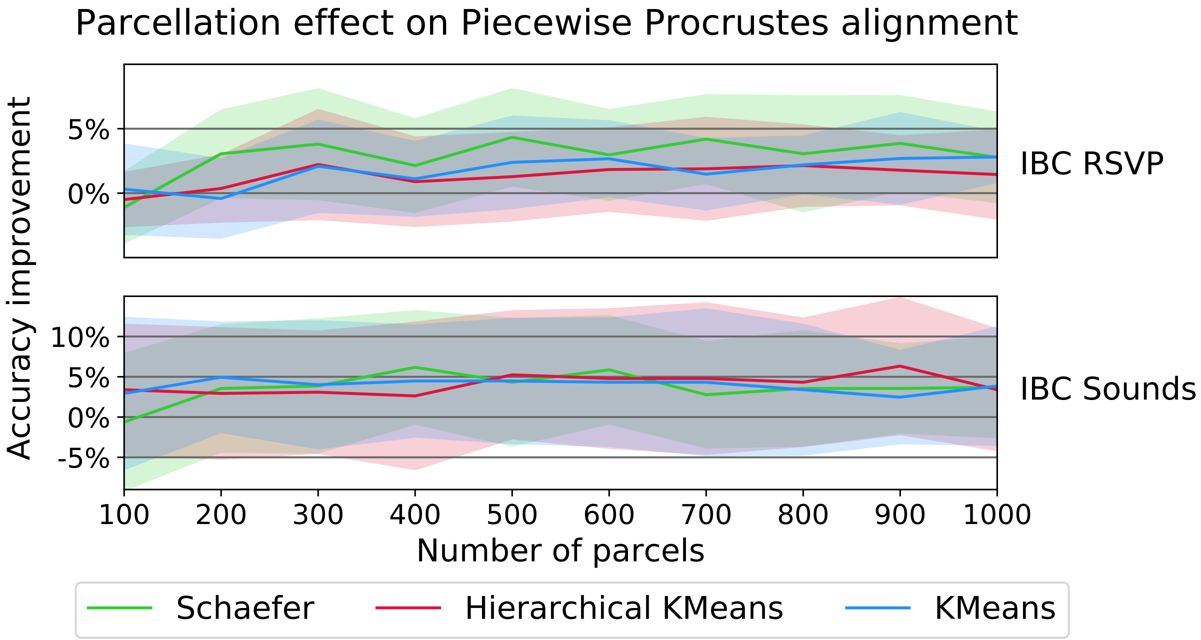


FIG. S2. **Effect of parcellation type, resolution on Piecewise Procrustes decoding accuracy improvement over anatomical alignment.** We consider the impact of parcellation type (the *a priori* Schaefer atlas or learned directly on the data with k-means or hierarchical k-means) and resolution (from 100 to 1000 parcels). Results are shown for the IBC RSVP and IBC Sounds decoding tasks. Each line represents the average accuracy improvement for piecewise Procruses over standard, anatomical-only alignment, and the confidence band represents the range of accuracy improvements seen across all IBC subjects. Accuracy improvements are calculated by subtracting anatomical-only inter-subject decoding accuracy scores for the same leave-one-subject-out cross-validation fold. We see that parcellation type and resolution show limited impact on accuracy gains.

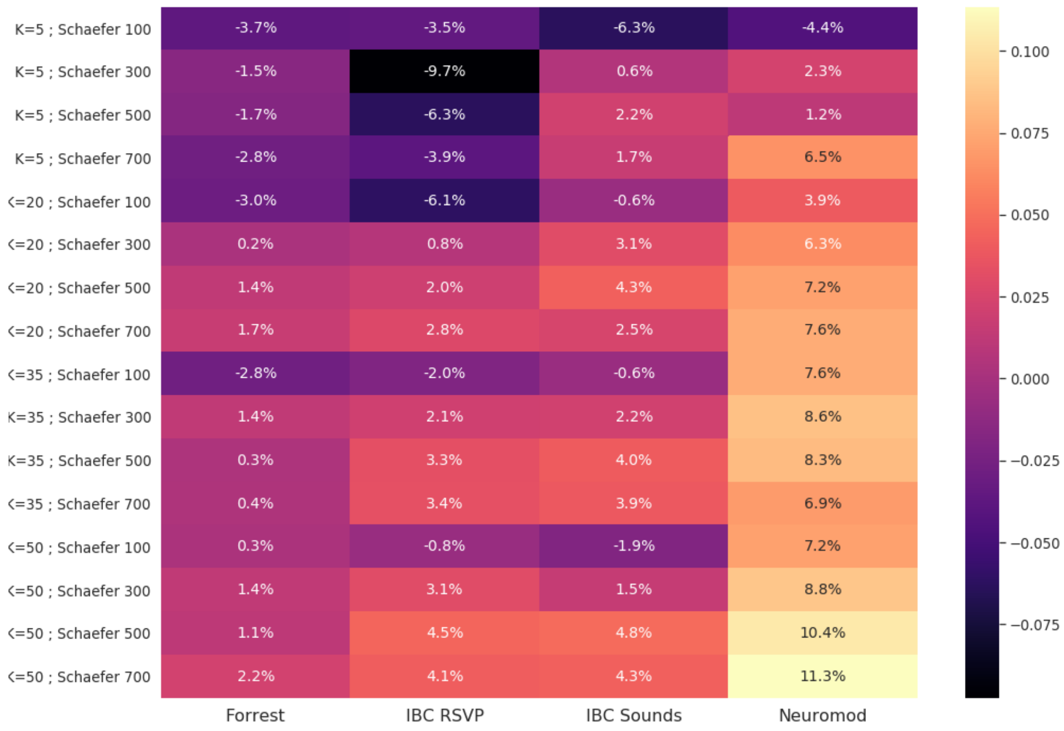


FIG. S3. **Grid search of Piecewise SRM hyperparameters impact on decoding accuracy across datasets** We considered a grid of 4 parcellations (Schaefer atlas at resolution 100, 300, 500 and 700) and 4 different values of $k$ (number of SRM components) for each model fitted on a parcel. We ran our inter-subject decoding pipeline on four inter-subject decoding task (in columns, among those used in the main benchmark). We report here the decoding accuracy improvement over anatomical baseline across datasets for each set of parameter (in line). Altough we didn't have the computational means to run an extensive grid-search, we can already conclude that high-resolution parcelations (and thus more fitted local SRMs) yield a higher decoding gain as long as they come with enough component. Decoding accuracy is also positively linked with $K$, probably up to a plateau that we did not clearly reached with our limited grid. For the main benchmark we retained the last line model ($K = 50$, Schaefer 700).

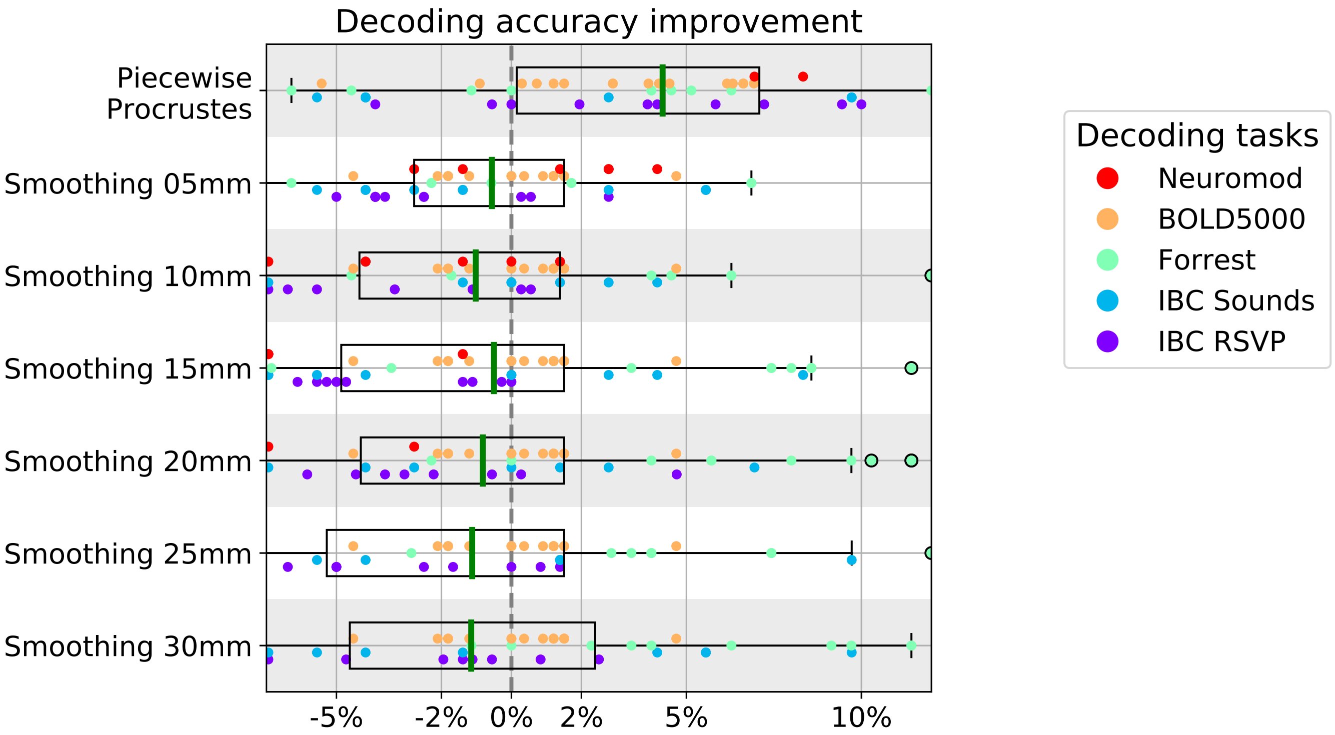


FIG. S4. **Decoding accuracy does not improve after Gaussian smoothing over anatomical alignment.** For six smoothing kernels, we show inter-subject decoding accuracy scores after subtracting anatomical-only inter-subject decoding accuracy for the same leave-one-subject-out cross-validation fold. Each dot represents a single subject, and subjects are colored according to their decoding task. We also show differences in decoding accuracy scores for the reference functional alignment method piecewise Procrustes, again as compared to anatomical-only alignment. Each box plot describes the distribution of values for that smoothing kernel or alignment method, and the green line indicates the median. We see that Gaussian smoothing does not show the same pattern of decoding accuracy differences as the reference functional alignment method.

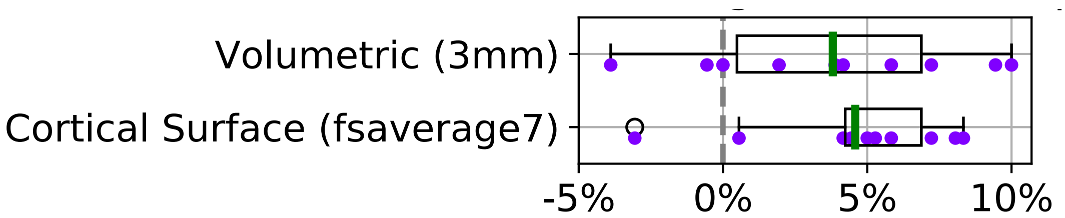


FIG. S5. **Comparing piecewise Procrustes accuracy improvements across volumetric and surface data representations.** For the IBC RSVP task, we compare piecewise Procrustes decoding accuracy scores to anatomical-only alignment. Each dot represents an IBC subject, where their difference score is calculated by subtracting the inter-subject decoding accuracy for anatomical-only alignment from the piecewise Procrustes alignment accuracy score for the same leave-one-subject-out cross-validation fold; i.e., where they are the left-out subject. We compare these difference scores as calculated using data in the volume (3mm resolution), to data on the high-resolution cortical surface (*fsaverage7*). Each box plot describes the distribution of values for that data representation, and the green line indicates the median. We see that the high-resolution surface representation yields a moderate gain of decoding accuracy, compared to 3mm isotropic volumetric representation.
